# Predicting gene expression using morphological cell responses to nanotopography

**DOI:** 10.1101/495879

**Authors:** Marie F.A. Cutiongco, Bjørn S. Jensen, Paul M. Reynolds, Nikolaj Gadegaard

## Abstract

Cells respond in complex ways to topographies, making it challenging to identify a direct relationship between surface topography and cell response. A key problem is the lack of informative representations of topographical parameters that translate directly into biological properties. Here, we present a platform to relate the effects of nanotopography on morphology to function. This platform utilizes the ‘morphome’, a multivariate dataset containing single cell measures of focal adhesions, the cytoskeleton, and chromatin. We demonstrate that nanotopography-induced changes in cell phenotype (both morphological and functional) are uniquely encoded by the morphome. The morphome was used to create a Bayesian linear regression model that robustly predicted changes in bone, cartilage, muscle and fibrous gene expression induced by nanotopography. Furthermore, the morphome effectively predicted nanotopography-induced gene expression within a complex co-culture microenvironment. The spatial, morphological and functional resolution of the morphome uncovered previously unknown effects of nanotopography on selectively altering cell-cell interaction and osteogenesis at the single cell level. Thus, the morphome confers the ability to quantify phenotype arising from cell-material interactions. Our new platform shows promise for rapidly assessing novel surface-patterned biomaterials for tissue regeneration and enables cell function-oriented exploration of new topographies.

## Introduction

Biomedical implants continue to be developed to improve patient outcomes. One way to enhance implant efficacy and tissue regeneration is to vary substrate texture with nanotopographies. Topographies at the cell-material interface are widely shown to direct cell behaviour: nanopillars change cell morphology^1^; nanogratings drastically alter lipid metabolism^2^, and pluripotent^3-5^ and multipotent cell differentiation^6^; nanogradients improve stem cell cardiomyogenic differentiation; and subtle changes to nanopit geometric arrangement switches human mesenchymal stem cells from multipotent to osteogenic fate^7-10^. Morphological responses to nanotopography are manifested through varying focal adhesion size^17-19^, orientation^19-21^, and composition^22,23^ and changes in actin contractility and nuclear deformation^24^.

A quantitative relationship exists between a material’s physicochemical structure and its biological activity. Rational drug design has long relied on molecule solubility, ionisation and lipophilicity to predict activity^11^. Protein engineering has similarly modelled protein-peptide interactions from protein structure^12^. Cell metabolic activity correlates with synthetic polymer composition, glass transition temperature, and water contact angle^13^. Meanwhile, bacterial attachment can be predicted from descriptors of secondary ionic hydrocarbon chains^14^. In contrast to active biomolecules, the mechanotransductive effects of topography on cell response do not intuitively relate to topography length scale, isotropy, geometry, and polarity. This limits the discovery of functional topography to the screening of libraries for hits using a single, representative cell type^15-18^. Among its limitations (which include cost, inefficiency and the sampling of a small topography space), this screening approach disregards the cell specificity of response to nanotopogrpahy. Thus, it is vitally important to develop a systematic method to capture cell phenotypes (both at morphological and functional levels) induced by topography.

Here, we demonstrate an image profiling-based platform that encompasses cell function and morphological response induced by nanotopography. Measurements of focal adhesions, actin cytoskeleton and chromatin, referred to here as the ‘morphome’, were obtained from single cells on nanotopographies. The morphome signature clearly reflected cell type, nanotopography and levels of gene expression at the single cell level. We used a Bayesian linear regression model to directly relate the morphological and phenotypic changes induced by nanotopography. Using the morphome as predictors and without prior knowledge about the nanotopography, the model robustly predicted gene expression induced by nanotopography. Precluding topography measurements (e.g. diameter, pitch of topography) in the model and the mechanosensitive response in all cell types highlights the broad applicability of this platform to many biomaterial and cell systems. In fact, a new morphome from a co-culture of osteoblastic and fibroblastic cells verified this by positively predicting level of osteogenesis induced by nanotopography. This is the first study to quantify morphological, functional and topographical relationships induced by cell-material interaction at a single cell level.

## Results

### Measuring morphological cell responses to topography: the morphome

We used nanopit topographies consisting of 120 nm diameter, 100 nm depth and with a 300 nm centre-to-centre distance in a square array (SQ)^7^, hexagonal array (HEX)^25-27^, and arranged with centre-to-centre distance offset from 300 nm by 50 nm in both x and y directions (NSQ array)^7,8^. An unpatterned (‘FLAT’) surface was used as a control (Figure 1A).

**Figure 1.**
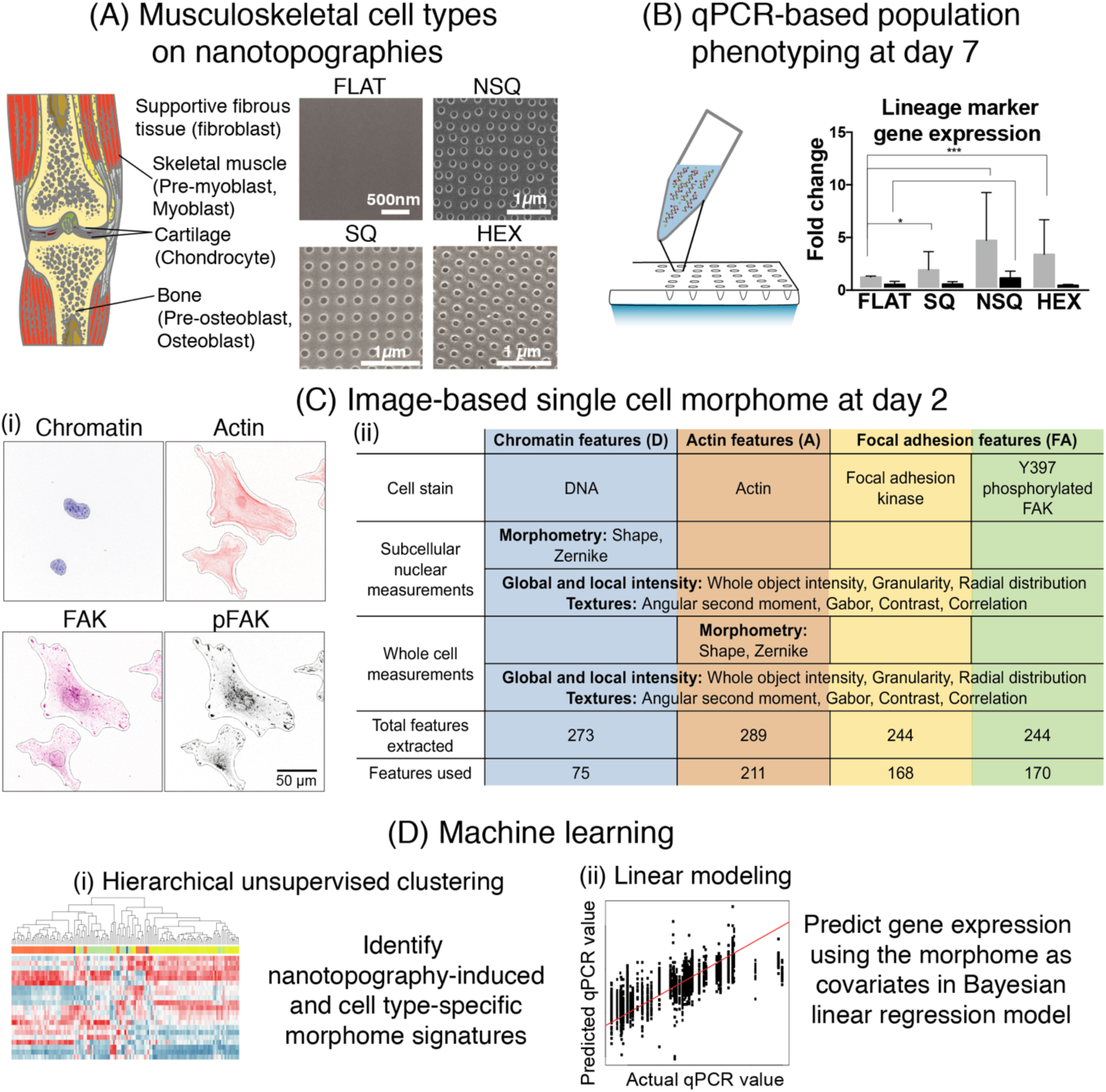
An overview of the morphome platform for relating cell function and morphology changed nanotopography. (A) Osteoblastic, myoblastic, chondroblastic and fibroblastic cell lines were ltured on nanotopographies to obtain 24 combinations of cell type and topography. Image of musculoskeletal system obtained from Servier Medical Art under creative commons licence 3.0. (B) day 7, lineage-specific gene expression induced by nanotopography was measured using population-based qPCR. (C) At day 2, various measures of chromatin (blue), actin (red), focal adhesion kinase (FAK, yellow) and phosphorylated FAK (pFAK, green) were obtained from images of single cells to create the ‘morphome’. (i) Representative images of cells stained against different cellular aspects. Black lines show the cell and nucleus outlines, from which morphology measurements were extracted. Cell and nucleus outlines were obtained from the actin and chromatin images, respectively, using CellProfiler. (ii) Cell morphology features extracted from 4 stains across single cells. (D) Machine learning for data-driven exploration and model building using the morphome. (i) Hierarchical clustering uncovered unique patterns that delineate cell type and nanotopography within the morphome. (ii) Bayesian linear regression created a predictive model that related the morphome to gene expression.

We employed cells of the musculoskeletal system due to the diverse responses of muscle, bone, cartilage and fibrous cell types to nanotopographies^27-29^. Mouse myoblasts, osteoblasts, chondrocytes, fibroblasts, and pre-osteoblast^30^and pre-myoblast^31^progenitors, were grown on nanotopographies. Responses from combinations of each cell type on all nanotopographies were measured, providing 24 unique combinations of cell type and nanotopography (Figure S1). Effects of nanotopography on conventional morphological characteristics such as cell and nuclear area, actin intensity, focal adhesion area and intensity, and mechanosensor nuclear translocation were evident (Figure S1-S4).

Quantitative polymerase chain reaction (qPCR) was then used to assess changes in lineage marker expression induced by nanotopography by day 7 (Figure 1B). At day 2, we performed image-based profiling (Figure 1C). From images of the chromatin and actin, the nucleus and the cell body, respectively, was robustly segmented using CellProfiler. Within these cellular features, we measured morphology (shape and geometry of different compartments), texture (spatial patterns of fluorescence), intensity (total fluorescence value), and radial distribution of intensity (measuring radial arrangement of fluorescence) of chromatin, actin, focal adhesion kinase (FAK) and phosphorylated FAK (pFAK) signals. The morphome consisted of 624 single cell measurements (“features”), with 75 chromatin and nuclear features, 211 actin and whole-cell features, 168 FAK features, and 170 pFAK features (Supplementary data). Machine learning was then applied on the morphome: (i) hierarchical clustering was used to uncover unique patterns of morphological features that distinguish cell type-specific responses on nanotopographies, and nanotopography-specific morphological changes induced within the cell; (ii) Bayesian linear regression was used to predict myogenic, osteogenic, chondrogenic and fibrogenic gene expression induced by nanotopography using the morphome as predictors (Figure 1D).

### The morphome depicts unique cell signatures induced by nanotopographies

Patterns of nanotopography-induced morphological changes were visible from the morphome (Figure 2A). Immediately apparent were large blocks of actin, FAK and pFAK measurements with similar values within a cell type on a specific nanotopography. These features correspond to increasingly complex measures of texture, granularity and radial intensity distribution for chromatin, actin, FAK and pFAK (Table S1). Frequency of pixel gray levels measure texture and homogeneity of pixels, with high values indicating coarseness. Granularity measures an object’s coarseness, with higher values indicating heterogeneity of pixel intensities and coarser texture. The Zernike coefficient measures the spatial arrangement of intensity as it resembles the increasingly complex Zernike polynomials (Figure 2A). The Zernike coefficient was used to measure both cell shape and radial distribution of fluorescence intensity of chromatin, actin, pFAK and FAK. Interestingly, higher order Zernike polynomials resemble the punctate shape and spatial distribution of focal adhesions^32^. This provides an integrative analysis of focal adhesions at the single cell level compared to traditional measures that define individual adhesion characteristics.

**Figure 2.**
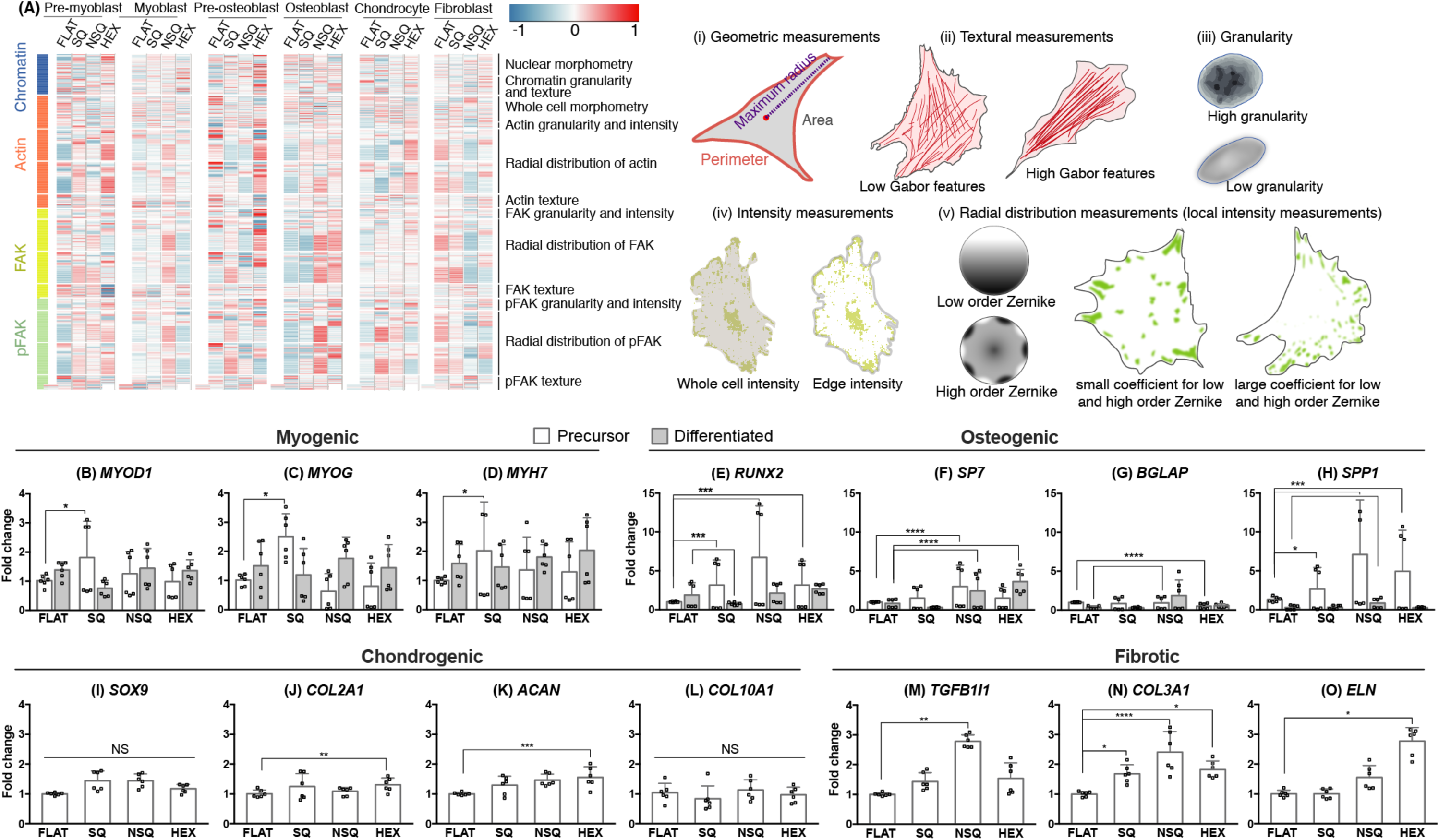
Unique responses of cells to nanotopography at single-cell and population levels. (A) Heat map of the morphome across cell types and nanotopographies. The morphome consisted of 624 features that quantitatively measure the cell and nucleus geometries, as well as chromatin, actin, FAK, pFAK characteristics within single cells. The colour and intensity of each tile represents the average value of the feature for a particular cell type and nanotopography combination. Each feature has an average of 0 and a standard deviation of 1 after normalization across all cell types. Schematic diagrams of representative morphome features are included in (i) to (v). (B-O) QPCR of musculoskeletal genes in response to nanotopographies. Measurement of (B-D) myogenic markers from pre-myoblasts and myoblasts, (E-H) osteogenic markers from pre-osteoblasts and osteoblasts, (I-L) chondrogenic markers from chondrocytes, (M-O) fibrotic markers from fibroblasts on nanotopographies. Gene expression is listed left to right in order of increasing maturity for the given cell lineage. QPCR measurements shown here were normalized to the reference gene and cell type on FLAT. QPCR measurements across all 24 combinations of cell type and topography are shown in Figure S5. All qPCR measurements are given as mean ± standard deviation from 2 independent experiments (n=6). Significance levels obtained from one-way ANOVA with Tukey’s post-hoc test for pairwise comparison. Significance levels were denoted by * (p<0.05), ** (p<0.01), *** (p<0.001), and **** (p<0.0001).

### Nanotopography induces cell type-specific gene expression changes

Gene expression was used to determine the effect of nanotopographies on cell function (Figure S5). We discuss here the changes induced by nanotopography on lineage-specific gene expression relevant to the cell type. Pre-myoblasts showed significantly higher expression of the early lineage marker *MYOD1*, and of the late markers *MYOG* and *MYH7* when cultured on SQ surfaces relative to FLAT surfaces (Figure 2B-2D). Both pre-osteoblasts and osteoblasts showed increased expression of early (*RUNX2, SP7*) and late (*BGLAP, SPP1*) osteogenic markers when cultured on NSQ relative to FLAT (Figure 2E-2H), in line with previous studies^7,8,10,33^. Chondrocytes cultured on HEX showed increased expression of *COL2A1* (early marker) and *ACAN* (late marker) compared to those cultured on FLAT (Figure 2I-2K). Meanwhile, fibroblasts showed increased expression of pathogenic fibrosis markers, *TGFB1I1, COL3A1* and *ELN*^*34*,^ on all nanotopographies compared with FLAT (Figure 2M-2O). In general, a cell type-specific response to nanotopography was observed. Each lineage was notably enhanced in a specific cell type and nanotopography combination: SQ stimulated the myoblast phenotype, NSQ enhanced the osteoblast phenotype, HEX stimulated the chondrocyte phenotype, while all surfaces except FLAT stimulated the fibrotic phenotype.

### Distinct nanotopographical responses of single cells are reflected in the morphome

A subset of the morphome, consisting of 185 features, varied significantly across cell types (Figure 3, Supplementary Data). For ease of visualization of a multivariate dataset, hierarchical clustering was employed to group morphome features of high similarity together and thus reveal morphological profiles across different nanotopographies. Before clustering, each morphome feature was mean centered and normalized, transforming each morphome feature to a relative scale with negative values denoting decrease and positive values denoting increase from the mean = 0. When taken entirely, hierarchical clustering of the morphome revealed distinct morphological profiles of all combinations of cell type and nanotopography (Figure S6). Our results were further distilled to hierarchical clustering of the morphome separated by cell type (Figure 3). Here, we highlight the morphome for cell type and nanotopography combinations that induced highest lineage-specific gene expression. When compared to FLAT, pre-myoblasts on SQ showed high average values of: focal adhesion textures (Figure 3A, cluster 2); pFAK radial distribution (cluster 4); nuclear morphometry; and chromatin textures (cluster 4). The morphome of pre-myoblasts cultured on SQ reflects the need for FAK phosphorylation and for its preferential localization at stress fiber edges, which is necessary for myotube differentiation^36,37^. In contrast, myoblasts on SQ showed a particularly high average value for chromatin granularity and nuclear morphometry (cluster 4), and near-zero values for radial distribution of actin and of focal adhesions (cluster 1). High chromatin granularity observed for both pre-myoblasts and myoblasts on SQ denotes chromatin heterogeneity and condensation and transcriptional activity, which is reportedly higher prior to myotube formation^38,39^.

**Figure 3.**
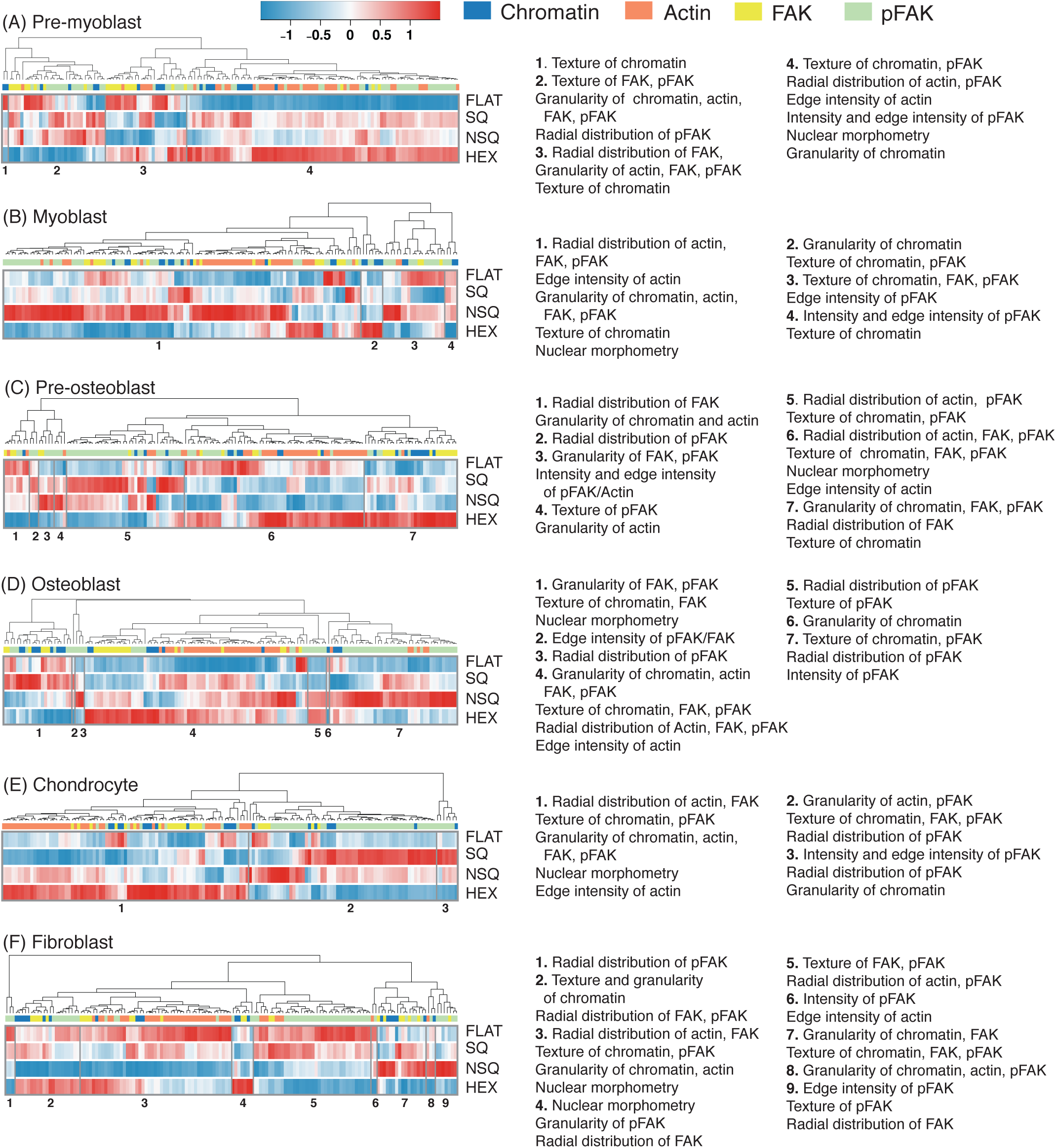
Morphome features hierarchically clustered by cell type revealed nanotopography-specific changes to cell morphology. (A-F) Hierarchical clustering of the morphome within each cell type. The data morphome from each cell type were clustered separately. Within the cell type-specific morphome, each morphome feature was normalized to have a mean= 0 and standard deviation = 1. The colour and intensity of each tile represents the average value of the feature. The morphome features analysed were comprised of 21 chromatin, 42 actin, 23 FAK and 60 pFAK features. Features included were changed significantly across topographies (p < 0.05 using one-way ANOVA). The total number of cells analyzed from two independent experiments were: (A) n=877 pre-myoblasts; (B) n=931 myoblasts; (C) n=644 pre-osteoblasts; (D) n=728 osteoblasts; (E) n=619 chondrocytes; and (F) n=1140 fibroblasts.

Pre-osteoblasts on SQ and NSQ had high average values for pFAK radial distribution, intensity, granularity and texture, and high average values for granularity of chromatin and actin (clusters 1-5). However, pre-osteoblasts on SQ had higher order pFAK and FAK radial distribution than on NSQ, which induced the highest expression of osteogenic markers. The morphome of pre-osteoblasts grown on NSQ featured radially variable actin that resemble bone cells, which have high contractility and actin stress fibers^40^.

For osteoblasts, the differences between the SQ and NSQ morphome were more prominent: NSQ induced lower average values of focal adhesion granularity, chromatin texture and nuclear morphometry (cluster 1), and higher average values for focal adhesion radial distribution (clusters 4-7) compared to SQ (Figure 3D). The osteoblast morphome on NSQ indicates that focal adhesions localize at regular intervals along the periphery, which is associated with osteogenesis^41^. Furthermore, changes in nuclear morphometry attributed to spreading after growth on stiff surfaces is also associated with osteogenic differentiation^42^.

Chondrocytes on HEX, which significantly increased chondrogenic marker gene expression relative to FLAT, showed high average values of radial distribution, texture and granularity of actin and FAK, high average nuclear morphometry, and low average values of pFAK and chromatin measurements (Figure 3E, clusters 1-3). These characteristics reflect the morphological changes (including reduced contractility and stress fiber formation, increased cell circularity, and decreased cell spreading^40^, low FAK phosphorylation^43^and poor focal adhesion formation^44^) of stem cells undergoing chondrogenesis.

The morphome of fibroblasts cultured on FLAT had high average values of both actin and focal adhesion measurements (Figure 3F, clusters 3 and 5). The highly uniform radial arrangement of focal adhesions and actin of cells on FLAT indicate reduced polarization and contractile morphology of fibroblasts activated to a fibrotic state^45^. Inflammation pathways are reportedly increased in fibroblasts on HEX^46^, inducing low adhesion that is reflected in low actin and focal adhesion radial distribution (clusters 3-9). Fibroblasts grown on NSQ and HEX showed low average values of focal adhesion and actin radial distribution but high values when grown on SQ (cluster 2-5).

Overall, the morphome reflected cell-type specific responses to nanotopography. This was highlighted by the dissimilarity of cell types with similar lineage or origin (e.g. pre-osteoblasts vs osteoblasts, Table 1). A multi-class logistic regression classifier was able to accurately distinguish 6 different cell types using the morphome (Figure S7). The radial arrangement of actin and focal adhesions were a critical distinction of the musculoskeletal cell types induced by nanotopography, while the arrangement of actin fibers into stress fibers or into cortical, circular bundles provided information on various cell states.

**Table 1.** Correlation coefficient of morphome features hierarchically clustered by cell type.

|  |  |  |  |  |  |  |  |
| --- | --- | --- | --- | --- | --- | --- | --- |
|  | All cell types | Pre-myoblast | Myoblast | Pre-osteoblast | Osteoblast | Chondrocyte | Fibroblast |
| All cell types | 1 |  |  |  |  |  |  |
| Pre-myoblast | 0.392 | 1 |  |  |  |  |  |
| Myoblast | 0.274 | 0.280 | 1 |  |  |  |  |
| Pre-osteoblast | 0.429 | 0.236 | 0.218 | 1 |  |  |  |
| Osteoblast | 0.410 | 0.201 | 0.280 | 0.354 | 1 |  |  |
| Chondrocyte | 0.540 | 0.164 | 0.374 | 0.393 | 0.397 | 1 |  |
| Fibroblast | 0.473 | 0.313 | 0.408 | 0.350 | 0.381 | 0.485 | 1 |

We also clustered the morphome based on nanotopography (Figure S8, Supplementary data). Patterns emerge in the morphome in direct response to nanotopography: NSQ induced high average values of pFAK radial distribution, texture and granularity; and HEX induced high average values of actin radial distribution. Correlation between the dendrograms confirm that the morphome clusters of different nanotopographies were dissimilar to each other (Table S2). The morphome enabled discrimination between 4 nanotopographies with 68% overall accuracy (87% classification rate for HEX) using multi-class logistic regression (Figure S9).

### The morphome predicts nanotopography induced gene expression changes

The Spearman rank correlation revealed that varying degrees of correlation exist between morphome features and gene expression (Figure S10). We hypothesized that the features of the morphomes would sufficiently encompass cell phenotypes induced by nanotopography. We utilized Bayesian linear regression to predict gene expression using the morphome features as predictors (for the explicit model definition, see Materials and Methods). A Bayesian linear regression model reflects uncertainty in the estimation of regression weights compared to point value estimates using maximum likelihood regression. Gene expression was modelled independently of each other, thereby creating 14 different equations with variable weighting of the morphome features. Importantly, the model was trained without any prior knowledge of topography type or parameters (e.g. nanodot diameter or center-to-center distance), instead relying on the morphome to encode this information.

The morphome clearly captured gene expression changes induced by nanotopography (Figure 4A). The heterogeneity inherent in single cells, usually uncaptured by population measurements of gene expression, are apparent in the variance of the predictions using the morphome and the model. The mean absolute error (MAE) for prediction of all genes was between 10% (for prediction of *MYOD1, MYOG* and *MYH7*) and 21% (for prediction of *COL3A1*, Table S3).

**Figure 4.**
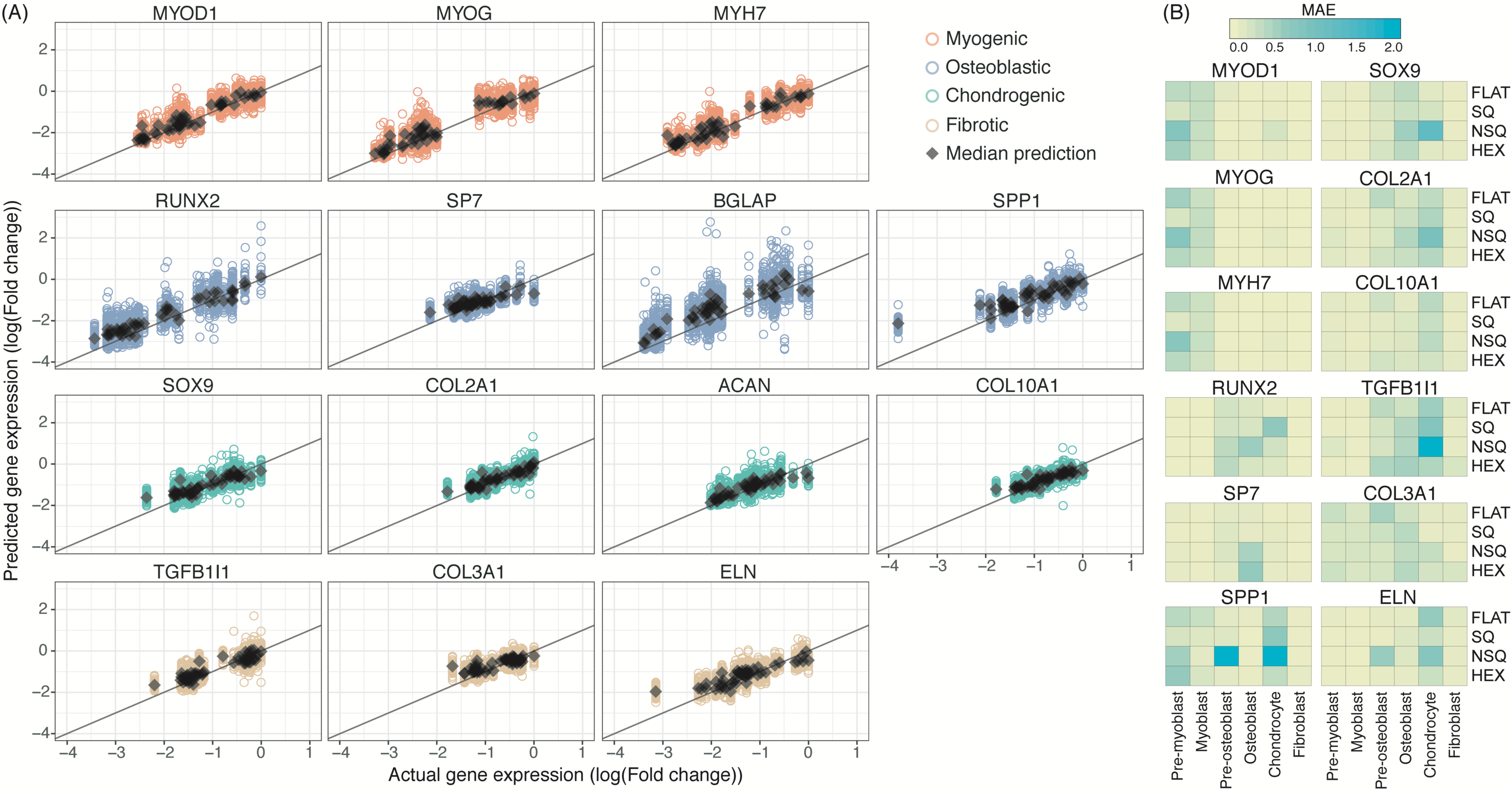
The morphome reliably predicts nanotopography-induced gene expression from specific cell type and topography combinations. (A) The morphome was used to train a Bayesian linear regression model that predicted myogenic, osteogenic, chondrogenic, and fibrotic gene expression. The expression of each gene (response) was trained against linear combinations of morphome features (predictors) and without any prior knowledge on topography parameters. The linear regression model was trained using 60% of the dataset and tested using 40% of the data. Scatterplots show actual and predicted gene expression values by using the test set as input to the model. Mean absolute error (MAE) was obtained by averaging the differences between actual and predicted qPCR values. (B) Testing the predictive power of the morphome by leave-one-out validation. To test the predictive power and bias of the morphome, the linear regression model was retrained after exclusion of one combination of cell type and topography. The excluded cell type and topography dataset was used for prediction, from which MAE was calculated. The tile position denotes the cell type and nanotopography combination that was excluded in the model and used for testing, while the color of each tile denotes the MAE.

The magnitude of the regression weight of each morphome feature reflects the contribution of the morphome feature in predicting gene expression. Across all 14 genes, pFAK activation, as indicated by pFAK/FAK integrated intensity ratio, consistently contributed to the prediction of all 14 genes. FAK texture and radial distribution, actin texture, and chromatin granularity features considerably contributed to prediction of gene expression (Figure S11, Supplementary data). pFAK was particularly important to the model due to its relevance in contractility induced by nanotopography^19^, fibrosis and scar tissue formation^48^, *in vitro* osteogenesis^49,50^, and chondrogenic maintenance^43^. Through the variability in the model, we obtain realistic expectations of single cell predictions from population level measurements

The sensitivity and predictive power of the morphome was verified by iteratively training a model without one combination of cell type and topography (Figure 4B). Drastic increases in MAE were observed when lineage-specific genes were predicted using models that excluded the particular cell type lineage being tested, regardless of nanotopography. The results are logically explained by the fact that the particular cell type contributes the most information to prediction by virtue of its lineage. Removal of the morphome of the particular cell type in question thus drastically reduces the amount of information in the model that uniquely determines that particular cell type. Model prediction after removal of particular nanotopographies showed higher consistency indicating that the generalizability of the morphome model towards predicting gene expression in response to nanotopographies outside of FLAT, SQ, NSQ and HEX.

### The morphome encompasses cell response to topography within a complex environment

We demonstrate the application of the linear regression model by predicting the outcome of pre-osteoblasts and fibroblasts co-cultured on nanotopographies. A new morphome was obtained from all cells on the entire nanotopography (Figure S12). This co-culture morphome was then used as input in the Bayesian linear regression model to predict gene expression (Figure S13).

For visualization, the sum of predicted osteogenic (*RUNX2, SP7, BGLAP* and *SPP1*) and fibrotic (*TGFB1I1, COL3* and *ELN*) genes was plotted against the spatial coordinates of the pre-osteoblasts and fibroblasts. Osteogenic gene expression was highest on NSQ, which also induced concentrated areas of highly expressing cells (Figure 5A). These areas might represent hotspots or nuclei of osteogenic paracrine signaling induced by the NSQ nanotopography^9^. In contrast, osteogenic gene expression was low and homogenous on the FLAT, SQ and HEX topographies. The uniformity of cell distribution across each nanotopography (Figure S12) eliminates the possibility of inadvertent cell clustering as the origin of gene expression changes.

**Figure 5.**
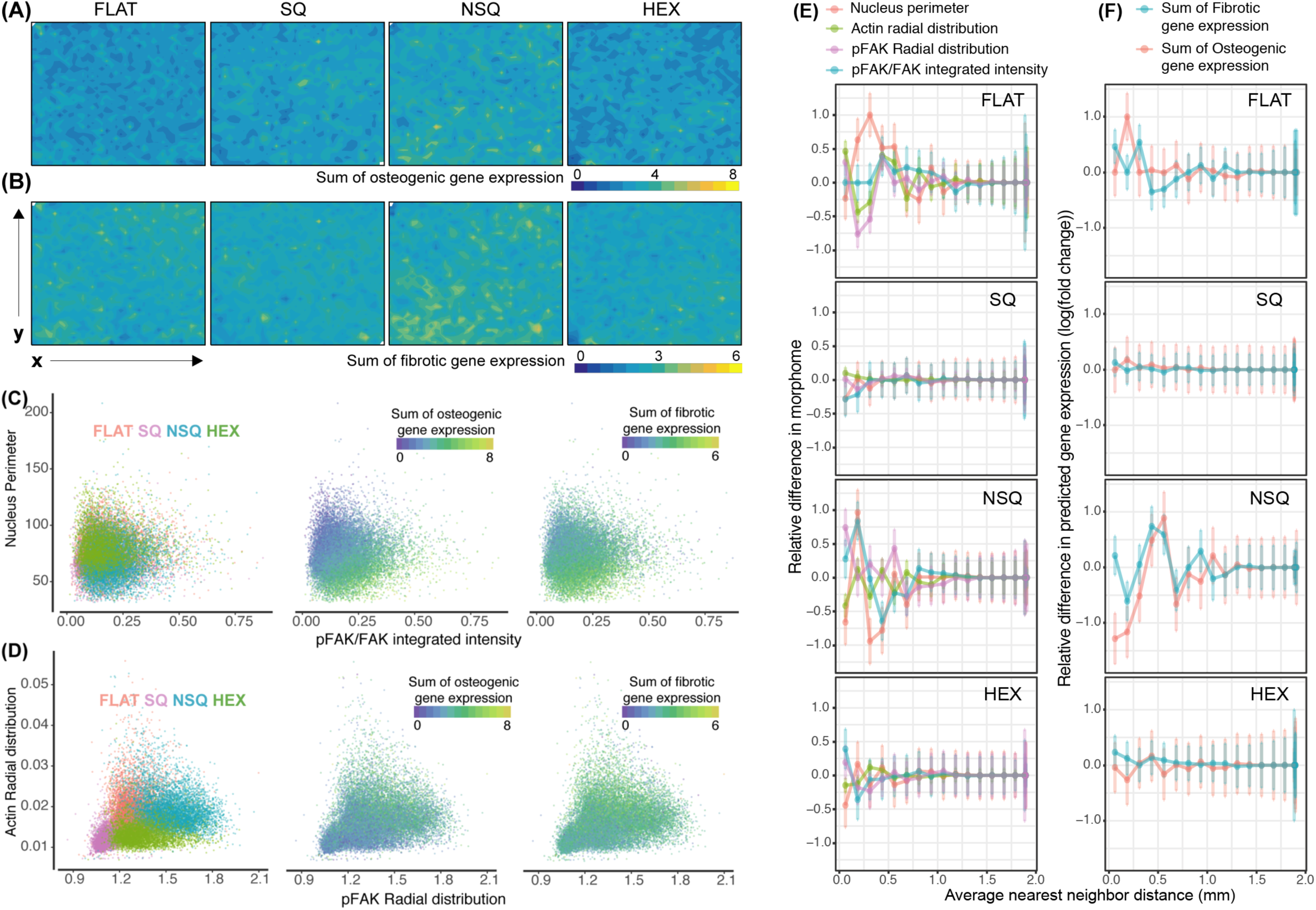
The morphome enables simulataneous analysis of morphological and functional changes induced by nanotopography at the single cell. (A-B) Predicted functional response of a pre-osteoblast and fibroblast co-culture to nanotopography. Contour plots show the sum of predicted (A) osteogenic (RUNX2, SP7, BGLAP, SPP1) and (B) fibrotic (TGFB1I1, COL3, ELN) gene expression for individual cells on FLAT, SQ, NSQ and HEX topographies. Pre-osteoblast and fibroblast cells were co-cultured on FLAT, SQ, NSQ and HEX nanotopographies for 2 days, and their morphome obtained from the entire grid. This new morphome was then used as input in the linear regression model (shown in Figure 4) to predict gene expression. X and Y axes of each contour plot shows are spatial coordinates on the nanotopogrpahy substrate. (C-D) Morphological and functional information at the single-cell level is provided by the morphome. Each dot in the plot denotes a single cell. (E-F) Cell-cell interaction altered by nanotopography. The average changes in (E) cell morphology and (F) gene expression between two cells separated by a given distance was measured and normalized to the maximum observed change. Data are presented as mean ± standard deviation and reported as a function of distance between two cells binned eveyr 125 µm. N ≥ 5000 cells per topography.

Fibrotic gene expression showed more spatial variability across nanotopographies but was also maximized on the NSQ nanotopography, and largely overlaps with the spatial pattern of osteogenic gene expression (Figure 5B). This is attributable to the synergistic interaction of osteoblasts and fibroblasts on osteogenic differentiation and mineralization^51^. The predicted effect of high osteogenic gene expression induced by NSQ was verified in the increased mineralization compared to FLAT at 28 days (Figure S14). Given that the morphome-based platform was created using single cell populations, the co-culture results indicate the capability to encode not just cell-material interaction but a concerted response arising from the cellular milieu.

For the first time, we gleaned new insights in single cell responses induced by nanotopography using the morphome. Due to the spatial, morphological and functional information at the single cell level, the morphome permits analysis of single cells similar to flow cytometry. As an example, we focused on correlating morphome features with high contribution to the predictive model and gene expression. Nanotopographies induced indistinct effects on either nuclear perimeter or pFAK activation, yet there was a clear gradient in predicted osteogenic gene expression (Figure 5C). As the nucleus became smaller and pFAK activation increased, both osteogenic and fibrotic gene expression increased. In contrast, nanotopographies exhibited clearly separable effects on actin and pFAK radial distribution, with cells on SQ showing the lowest values (Figure 5D). These particular changes in cell morphology correlated more with osteogenic gene expression than fibrotic gene expression, which lacked clear separation by nanotopography. From the morphome, we found strong evidence to implicate changes in nucleus shape and pFAK activation induced by nanotopography as drivers of osteogenesis.

The morphome additionally uncovered effects of nanotopography on cell-cell interaction at the morphological and functional levels (Figure 5E, 5F). On average, the effect of FLAT on cell-cell interaction and the resulting cell morphology extended up to 1 mm (Figure 5E), yet gene expression changed maximally at a separation distance of only 250-375 µm between neighboring cells (Figure 5F). In contrast, the average effect of SQ and HEX on cell-cell interaction and morphology were minute and apparent only at short cell-cell separation distances of 250 µm and 500 µm, respectively. A predominantly negative effect on pFAK activation was observed between neighboring cells grown on either SQ or HEX. This suppresive effect of nanotopography on cell-cell interaction correlate strongly with the homogeneity in gene expression induced by SQ and HEX (Figure 5F). On the contrary, long-range effects between cells on NSQ were distinct (Figure 5E). In contrast to FLAT, neighboring cells on NSQ separated by 1 mm or less selectively exhibited drastic changes only in nucleus perimeter and pFAK activation. The long-range effects of cell-cell interaction observed in NSQ were clearly manifested in gene expression (Figure 5F). In fact, NSQ showed a critical distance of 500 - 625 µm between neighboring cells where fibrotic and osteogenic gene expression were maximally changed. In summary, we observed a clear augmentation in cell-cell interaction induced by NSQ compared to FLAT, SQ and HEX. This long-range interaction between neighboring cells on NSQ separated by 1 mm or less drove changes in pFAK activation and osteogenesis.

## Discussion

In this study, we present a system that robustly encodes nanotopography parameters into functional cell specific output. The morphome, which in this study is the collective morphological measurements of chromatin, actin and focal adhesions within single cells, were found to manifest nanotopography-induced changes in cell morphology and gene expression. The information encoded in the morphome underpinned the performance of a Bayesian linear regression model for predicting gene expression. Using the morphome, we gleaned new biological insights resulting from nanotopographical perturbation of the cell microenvironment.

Focal adhesion complexes are the primary mechanosensing machinery of the cell that respond to the biophysical microenvironment. Focal adhesion assembly or confinement induced by nanotopography allows spontaneous actin cytoskeleton assembly, nucleus deformation, and cell fate determination^24,52,53^. Cells also change and control their microenvironment, thereby dictating focal adhesion and cellular characteristics to varying degrees^54,55^. Focal adhesion growth is particularly significant in sensing nanoscale changes in ligand arrangement, as this supports force redistribution throughout the cell ^56^. This reciprocity between the nanotopographical microenvironment, focal adhesions, cytoskeleton and chromatin, and gene expression underline the platform presented in this study. Our results demonstrate that cell morphological and functional response to nanotopography emerge from the morphome. The morphome-based platform reported here offers two unique advantages. First, the design parameters of the nanotopographies were explicitly excluded in predicting gene expression. To use structural information from topography (e.g. depth, diameter, pitch, geometry, isotropy, roughness) considerably limits the dataset to only a handful of descriptors. As opposed to polymer-based biomaterials that contain easily interpretable physicochemical properties derived from chemical structure, the effects of topography parameters on hydrophobicity, serum adsorption and cell attachment are less well understood and quantified. Instead, the information contained in the morphome encompass both topography and gene expression. Furthermore, by creating a general model that is independent of topographical parameters, this approach can be easily applied to predict cell phenotype induced by new nanotopographies, given a set of images of cells grown on them. Second, in this approach, gene expression-based measures of cell phenotype determined the specialized function of cells. In contrast to our system, many quantitative structure and function relationship studies of polymeric biomaterials have focused only on adhesion and metabolic activity^13,14^. Since gene expression is highly scalable, the linear regression model can be adapted to predict other cell behaviours. This also ensures that the morphome supports a function-focused exploration of the topography space and the rational design of topographies, as opposed to the currently used trial-and-error screening approach.

Our data reveals that the morphome can also manifest nuanced information within the cellular lineage commitment timeline. The morphome easily captured higher levels of gene expression in cells farther along lineage commitment compared to precursor cells. Additionally, our co-culture experiments robustly predicted the osteogenic properties induced by NSQ. The elevated level of predicted osteogenic gene expression was supported by high mineralization on NSQ after 28 days of culture. Our results suggest that the morphome can manifest cellular changes induced by nanotopography, as well as changes driven by chemical or paracrine cues. This property of the morphome can be exploited to predict cell behavior in complex microenvironmental settings.

The co-culture experiment also demonstrated that the morphome dataset encompasses, at high resolution, structural, functional and spatial information. Indeed, we took advantage of this informationally rich dataset to uncover enhancement of cell-cell interaction (from micron to millimeter range) resulting from a subtle change in nanotopography order. SQ and HEX, both of which present an ordered interface to the cell, suppressed cell-cell interaction at 250 – 500 µm. Meanwhile, cell-cell interaction was apparent at long distances of 1 mm on FLAT and NSQ, which preferentially varied pFAK activation between neighboring cells. This result presents a new mechanism for nanotopography induced-cell behavior.

Clearly, morphome capture is crucial to the ability of the linear regression model to predict nanotopography induced gene expression. While population-level measures of gene expression strongly indicate cell function, they introduce a measure of uncertainty and biological variability into the linear regression model. Thus, a one-to-one relationship between the morphome and cell function is essential to develop. Non-destructive microscopic and molecular tools^57^that combine spatial and structural information from the morphome with single-cell functional assays are vitally important for establishing quantitative topography structure- and cell-function relationships using the morphome. However, the use of routine methods, such as high-content imaging and qPCR, permits any lab to measure the morphome and to model it against the gene expression in question.

By generating a multivariate morphome dataset and combining it with machine learning, we have created a powerful platform for relating topography structure to gene expression. The predictive power of the Bayesian linear regression model we have developed easily lends to sequential experimental design by exploiting uncertainty and variability within the model^58,59^. Combined with bench-top lithographic techniques^60^and *in silico* simulation of morphological response to nanotopography^61^, we envision a completely closed-loop system that enables functionally-oriented exploration of new topographies.

## Materials and methods

### Polycarbonate surfaces with nanotopography

Surfaces patterned with 120 nm diameter and 100 nm depth nanopits were fabricated on polycarbonate using injection molding^62^. The following nanotopographies were used: surfaces without nanopits (FLAT); nanopits in a square array with 300 nm center-to-center spacing (SQ); nanopits in a square array with approximately 300 nm center-to-center spacing distorted by 50 nm in both x and y directions (NSQ); nanopits in a hexagonal array with 300 nm center-to-center spacing (HEX). Samples were cleaned in 70% ethanol and dried before treating with O_2_ plasma at 120 W for 1.5 mins. Samples were sterilized using UV light in biological safety cabinet for at least 20 mins before cell seeding.

### Cell culture

Mouse fibroblast cell line NIH3T3 (ATCC) was cultured in reduced sodium bicarbonate content (1.5 g/L) Dulbecco’s modified eagle’s medium with (DMEM) supplemented with L-glutamate (2mM), 10% bovine calf serum, and 1% penicillin-streptomycin. Mouse C2C12 myoblasts (ATCC) were cultured in DMEM with 20% FBS and 1% penicillin-streptomycin, and differentiated into skeletal muscle using DMEM supplemented with 2% horse serum and 1% penicillin streptomycin. Mouse chondrocytes were cultured in minimum essential medium alpha (MEM*α*) with nucleosides, ascorbic acid, glutamate, sodium pyruvate supplemented with 10% FBS and 1% penicillin-streptomycin. Mouse MC3T3 cells (ATCC) were cultured in MEM*α* with nucleosides and L-glutamine without ascorbic acid and supplemented with 10% FBS and 1% penicillin-streptomycin. To differentiate MC3T3 into osteoblasts, MC3T3 media was supplemented with 10 nM dexamethasone, 50 µg/ml ascorbic acid and 10 mM β-glycerophosphate^63^. Lineage committed progenitor cells, referred to here as pre-osteoblasts and pre-myoblasts, were also included in the study to mimic the osteogenic and myogenic regeneration in the adult tissue^30,31^.

### Cell seeding

Cells were harvested from flasks using trypsin in versene buffer and spun down at 400 x g for 5 minutes. NIH3T3 and MC3T3 cells were resuspended in complete media and seeded at 4000 cells/cm^2^. Chondrocytes and C2C12 were seeded at 2500 cells/cm^2^. Cells were seeded at different densities to ensure single cells at approximately 30% confluency on each surface after 2 days culture. To ensure homogeneity of seeding, cells were seeded using a cell seeder that controlled fluid flow. For co-culture studies, MC3T3 and NIH3T3 cells were simultaneously seeded at 2000 cells/cm^2^ per cell type in MC3T3 growth media. All cells were grown for 2 days on nanotopographies before fixation for immunofluorescence staining.

### Quantitative polymerase chain reaction (qPCR)

After 6 days, total RNA was obtained from lysed cells according to manufacturer’s instructions (Promega ReliaPrep Cell Miniprep kit). Gene expression was measured directly from 5 ng RNA using a one-step RT PCR kit with SYBR dye (PrimerDesign). qPCR was run on the BioRad CFX96 platform. Relative gene expression was normalized to the 18S ribosomal RNA reference gene. A list of the forward and reverse primers used to study different mouse genes is given in Table S4. Bar charts for qPCR data were obtained using GraphPad Prism (v7.0). One-way ANOVA with Tukey’s post hoc test for multiple comparisons was performed to determine the effect of nanotopography on gene expression compared with FLAT. Statistical significance was considered at p < 0.05.

### Immunofluorescence staining

After 2 days, cells on surfaces were fixed with 4% paraformaldehyde solution in phosphate buffered saline (PBS) at 4°C for 15 minutes. Fixed cells were then permeabilized and blocked with 10% goat serum and 2% bovine serum albumin in PBS for 1 hour at room temperature. Surfaces were stained with the following primary antibodies overnight at 4°C: pFAK Y397 (Abcam 39967, 1:400) and FAK (ThermoScientific 396500, 1:400). Afterwards, Alexa Fluor-conjugated secondary antibodies (ThermoScientific, 1:500) against the host species of the primary antibody were used. Alexa Fluor 549 conjugated phalloidin (ThermoScientific, 1:200) were used to structuralize the actin cytoskeleton. Cells were also stained with DAPI (ThermoScientific) to visualize chromatin inside the nuclei. DAPI was previously reported contribute textural information as a means of alternatively representing chromatin^64^. All surfaces were mounted on 0.17 µm thick Glass coverslips with ProLong mounting medium (ThermoScientific) and dried overnight at 4°C before imaging.

### Image acquisition and feature extraction

For single-cell population studies, monochrome images of each fluorophore were obtained at 40X magnification (numeric aperture 1.3) using the EVOS FL1 System (ThermoScientific). For co-culture studies, the entire nanotopography field was imaged and stitched through an automated microscope (EVOS FL2 Auto) with a 40X magnification. All images from one cell type were obtained using the same camera and light settings. Afterwards, image processing and feature extraction were perfomed using CellProfiler^65^(v2.4.0, The Broad Institute). Image processing, including illumination correction and channel alignment, was performed across each independent experiment and each cell type^66^. Nucleus and cell body were segmented from each cell in each image, allowing single cell analysis of shape or morphometric measurements, total and local intensities and textures from chromatin, actin, pFAK and FAK.

### Multivariate analysis

The morphome initially consisted of a total of 1050 measurements obtained from single cells. Morphome measurements from single-cell populations were combined and were scaled by subtracting the mean and normalizing by the standard deviation of the entire data set. Morphome data from co-culture studies were similarly scaled and normalized using the mean and standard deviation from the initial dataset consisting of homogeneous cell populations. Prior to machine learning, features with zero variance within each batch (e.g. Zernike Phase measurements) were removed from the data set. A Pearson correlation method at significance level 90% was used to remove features with correlation higher than 0.9 without significantly reducing total data variance (Figure S15) using the *KMDA* (v1.0) package for R^67^. After pre-processing, 624 morphome features were used in the study.

### Hierarchical clustering

To determine the features that were significantly varied across nanotopography, a one-way ANOVA with Tukey’s post hoc test for multiple comparisons was performed (KNIME v3.3.0). Prior to clustering, each morphome feature was mean centered and normalized to the standard deviation. An agglomerative hierarchical clustering algorithm was performed using a Euclidean distance metric and an average linkage method for cluster linkage using *gplots* (v3.0.1) package^68^. Membership of each morphome in a cluster was obtained from silhouette analysis using the *cluster* (v2.0.7-1) package^69^. Hierarchical clustering was visually represented with a dendrogram, and a heatmap with color intensity corresponding to the average values of the morphome features. Dendrogram correlation was performed using the *corrplot* package^70^.

### Bayesian linear log-Normal regression

Only morphome features with an absolute Spearman correlation coefficient ≥ 0.7 against all examined expression markers were used in the linear regression model (Supplementary data). The linear regression model used 243 morphome features, containing 22 nuclear morphometry and chromatin, 71 actin, 75 FAK and 75 pFAK measurements, as predictors of the model. Each replicate of the qPCR data was propagated across each replicate of the single-cell morphome data. For each gene analysed, data were rescaled from 0 to 1 by normalizing to the maximum gene expression.

Linear regression was performed as a simple approximation of the relationship between the morphome and myogenic, osteogenic, chondrogenic and fibrotic gene expression. Established Bayesian inference methods were used to determine the probabilities of observing gene expression with a given morphome set. We consider a linear model where expression of one gene (response *y*) was predicted through a linear combination of the morphome features (predictors *X*) transformed by the inverse identity link function. We assume that *y* follows a log Normal distribution parametrized by the mean *µ* and variance 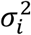:

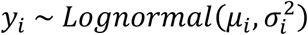

And that *µ* is a linear function of *X* parametrized by *β*:

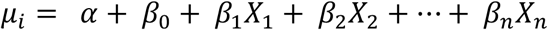

All model parameters *β* were assumed *a priori* to come from a normal distribution, parametrized by mean and standard deviation:

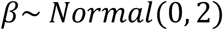

Each gene was trained independently resulting in 14 different linear regression equations. A 60%-40% training and test split for Bayesian linear regression was performed randomly and with stratification using the *caret* (v6.0-81) package for R^71^. The Bayesian linear model was created using the *brms* (v2.5.0) package for R^72^, which utilizes the Hamiltonian Markov Chain Monte Carlo sampler for estimation of the posterior distribution of *β*. Bayesian linear modelling was carried out using with 1000 warm-up iterations and 1000 sampling iterations within each chain for 3 independent chains. All models were confirmed to converge to the equilibrium distribution by confirming potential scale reduction statistic 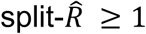, effective sample size was smaller than total sample size and low autocorrelation. Predicted gene expression was performed by using the test set or the morphome obtained from the co-culture study as input to the linear model. Predicted values were averaged across 50 draws from the posterior distribution. The magnitude of the average values of parameters *β* indicated feature importance as it effectively weighted the contribution of each morphome feature in predicting gene expression.

To determine the predictive power of the morphome, a specific combination of cell type, topography and replicate were iteratively omitted and the remaining dataset was used to refit new models. Thus, 576 additional models were created to test 24 different cell type combinations across 12 genes and 2 replicates. The predictive quality of the models was assessed by predicting the expression of all 14 genes from the omitted cell type, topography and replicate dataset. We report the mean absolute error (MAE) of qPCR prediction for each cell type and topography combination averaged across 2 replicates. MAE was calculated as the average across all absolute differences between predicted and actual gene expression.

### Analysis of cell-cell interaction

The co-culture morphome was used to predict gene expression at the single cell level. The dataset was then used to determine changes in gene expression and morphome between neighboring cells of a given distance. Changes in cell morphology and gene expression was performed for each cell against all other cells. Distances between cells were binned to calculate average value of changes in cell morphology and gene expression at 125 µm intervals. The changes in cell behavior between two cells was normalized to the maximum value of change observed.

### Statistics, visualization and software

Statistical analysis and machine learning were performed using statistical software R (v3.4.3) and its graphical interface RStudio (v1.0)^73^. Scatterplots, boxplots and histograms were generated using *ggplot2* (v3.1.0) in R^74^. Interpolation of x and y coordinates and sum of predicted osteogenic or fibrotic gene expression was performed using bivariate interpolation of a regularly gridded dataset using *akima* (v0.6-2) package^75^. Afterwards, the contour plots were created using the *fitted.contour* function in the base package of R, with the nuclear centroid position used as spatial coordinates of the cell. Barcharts and one-way ANOVA analysis of qPCR values were obtained using GraphPad Prism (v7.0a).

## Supporting information

Supplementary Data - pairwise comparisons

Supplementary Data - Weights of morphome feature in linear regression

Supplementary Data - Morphome features varied across cell type

Supplementary Data - Morphome features varied across topography

Supplementary Data - Morphome features

## Data availability

The data that support the findings of the study are available from the corresponding author upon reasonable request.

## Code availability

The statistical models proposed and evaluated in this paper is realised using standard packages in R. The code is available from the corresponding author upon reasonable request.

## Acknowledgements

We acknowledge ERC funding through FAKIR 648892 Consolidator Award. MFAC is financially supported by the University of Glasgow MG Dunlop Bequest, College of Science and Engineering Scholarship, and FAKIR consolidator award. We acknowledge the James Watt Nanofabrication Centre for fabrication work, and Steen Lillelund for initiating the machine learning work. We thank Julie Russell for her contribution to the qPCR, Carmen Huesa for providing the primary cartilage cells and Rachel Love for the injection moulding of nanopatterned polycarbonate surfaces.

## Author contributions

MFAC, PMR and NG designed the biological experiments. MFAC and BSJ designed the machine learning analysis. MFAC carried out imaging, image analysis, qPCR, machine learning. PMR fabricated and characterized the nanotopographical surfaces. MFAC, BSJ, PMR and NG wrote the manuscript. All authors have read and approved the manuscript before submission.

