## Supplemental Data 1 for "Predicting gene expression using morphological cell responses to nanotopography"

### Supplementary tables

**Table S1. Descriptors/features measured by various modules on CellProfiler.** Adapted from the CellProfiler manual[1], Kumar et al.[2], Vega et al.[3], and Huang et al[4].

| | | |
|---|---|---|
| **Area and Shape** | | |
| Area | Total number of pixels in each object. | 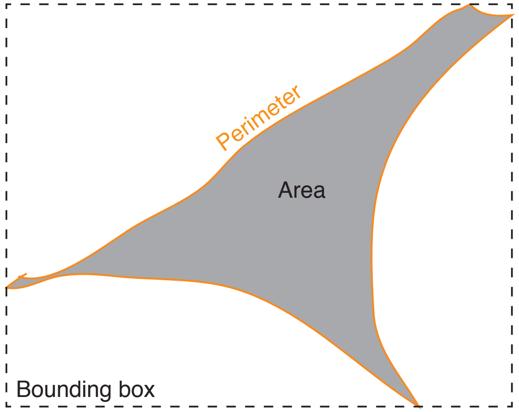 |
| Perimeter | Total number of pixels on the boundary of each object. | |
| Form Factor | Ratio of the area to the perimeter of each object. Ratio of 1 is for a perfect circle. | |
| Extent | Given the smallest bounding box that can enclose the object, the extent is the ratio of the area of the object by the area of the bounding box. Cells that have fewer protrusions have larger extent values. | |
| Eccentricity | Ratio of the distance between two foci and the major axis of the ellipse that | |

| | | |
|---|---|---|
| | bounds the cell. Ratio approaching 1 indicates a more elliptical shape. | 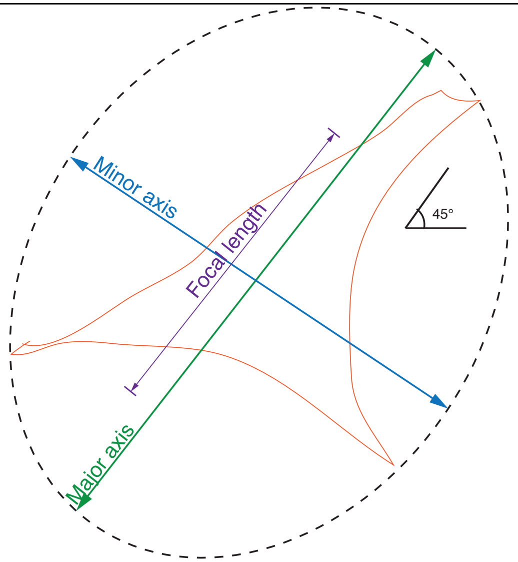  |
| Major Axis Length | Length of the major axis of the ellipse that bounds the cell given in pixels. | |
| Minor axis length | Length of the minor axis of the ellipse that bounds the cell given in pixels. | |
| Orientation | Angle between -90 and 90 degrees that indicates the orientation of the major axis of the cell with respect to the horizontal. | |
| Maximum Radius | The greatest distance between any pixel inside the object to the nearest pixel outside of the object. | 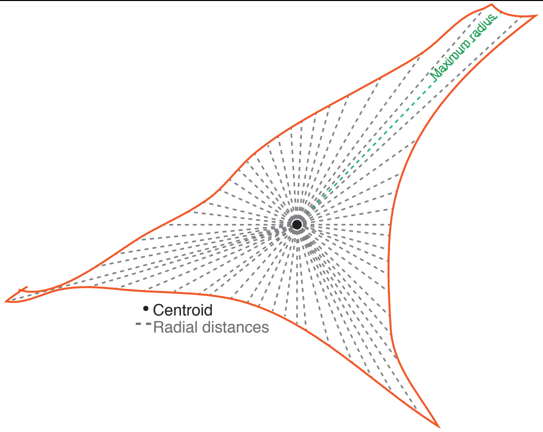  |
| Compactness | Ratio of the variance in the radial distance from the centroid and the area of the object. | |
| Maximum Feret Diameter | The maximum distance between two parallel lines tangent to opposite sides of the object. | |

| | | |
|---|---|---|
| Minimum Feret Diameter | The minimum distance between two parallel lines tangent to opposite sides of the object. | 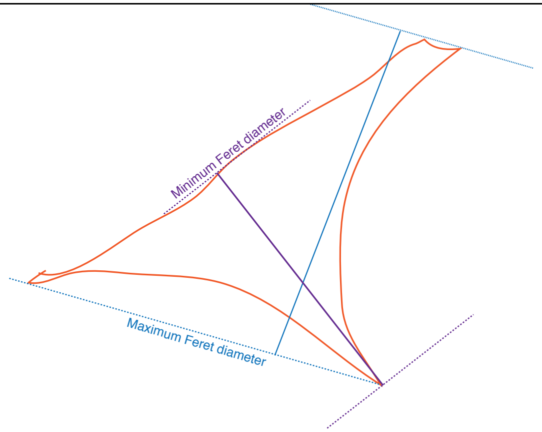 |
| Solidity | Measure of how many holes or concave boundaries in each object. Measured as area/convex hull area. The convex hull area is fitted onto the object by drawing a straight line across straight or concave areas of the cell but expanding outward along object protrusions. Ratio of 1 indicates an object without any concavities or indentations, while a ratio less than 1 indicates an object with holes or an irregular boundary. Ratio of Area to convex hull area. | 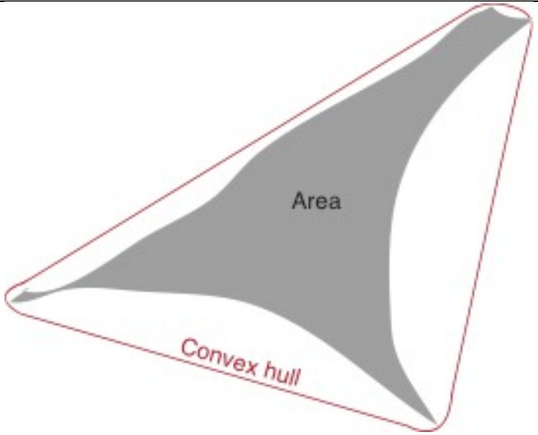 |
| Zernike shape features | Set of polynomial coefficients used to describe cell shape with increasing detail. The smallest circle that encloses the object is used to calculate Zernike features. Cells that are more closely related to the Zernike polynomial in question are reflected in a higher value. | 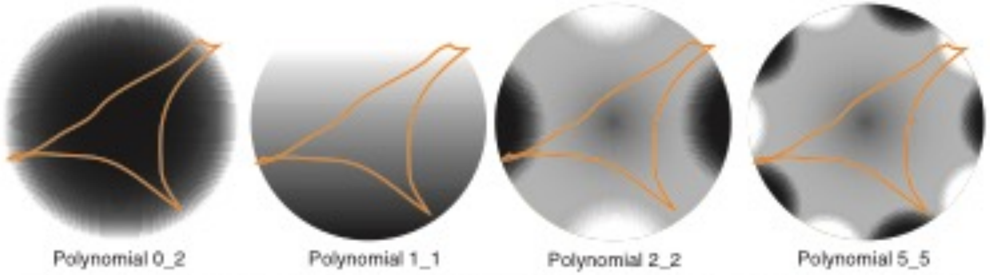<br>Cell is low in Zernike coefficients for polynomials 0_2 and 2_2 but high for polynomial 1_1 and 5_5 |

| Intensity | | |
|---|---|---|
| Integrated Intensity | Total intensity values of pixels within a specified object. (e.g. measurement of total intensity of focal adhesions within a single cell) | 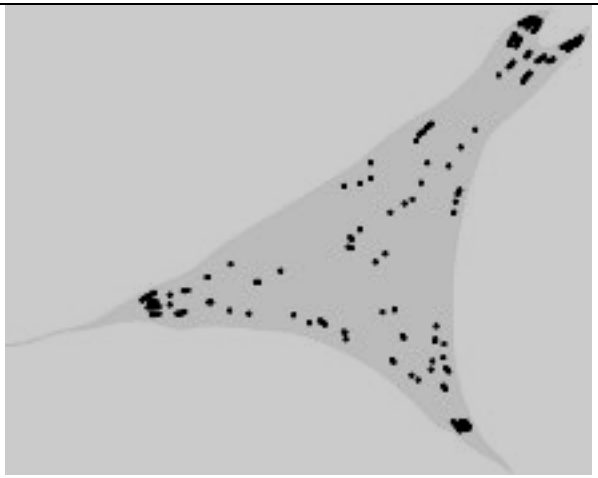 |
| Integrated Edge Intensity | The sum of pixel intensities along the perimeter of the object. (e.g. measurement of total intensity of focal adhesions on the perimeter of the object) | 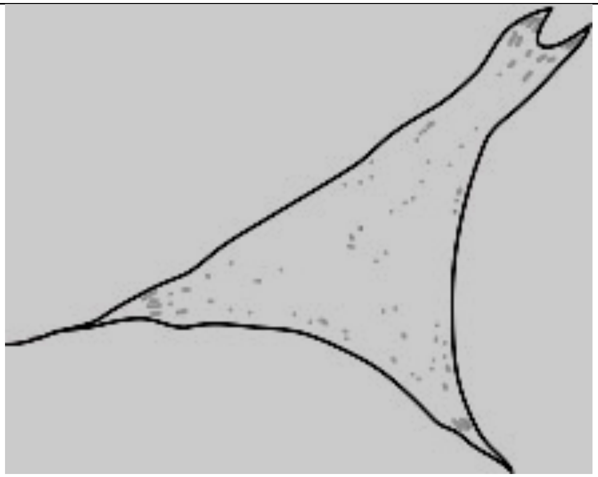 |
| Fraction at Distance (FracAtD) | The ratio of the stain in object to the total stain within a given radius (measured from the cell nucleus). | |

<table>
<tr><td>Mean Fractional Intensity (MeanFrac)</td><td>The mean fractional intensity at a specified radius (with the center defined as the nucleus).</td><td rowspan="2">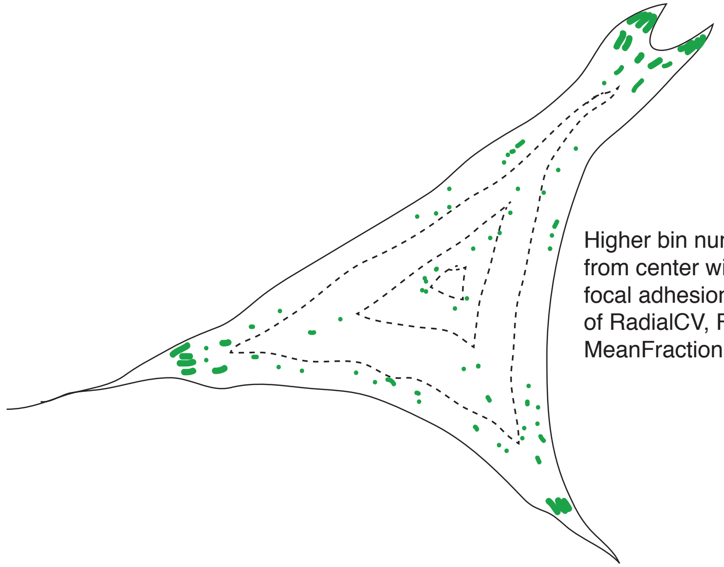<br>Higher bin numbers that are farther from center will high concentration of focal adhesions and have higher values of RadialCV, FractionAtDistance and MeanFractionalIntensity</td></tr>
<tr><td>Radial coefficient of variation (RadialCV)</td><td>Coefficient of variation of intensity within a ring (with the center defined as the nucleus).</td></tr>
<tr><td>Zernike Magnitude</td><td>Characterizes the distribution of intensity across an object from its center described by Zernike polynomials. Higher Zernike polynomials describe the object with higher detail. Values obtained are correlation coefficients between the intensity distribution within the object and the given Zernike polynomial.</td><td rowspan="2">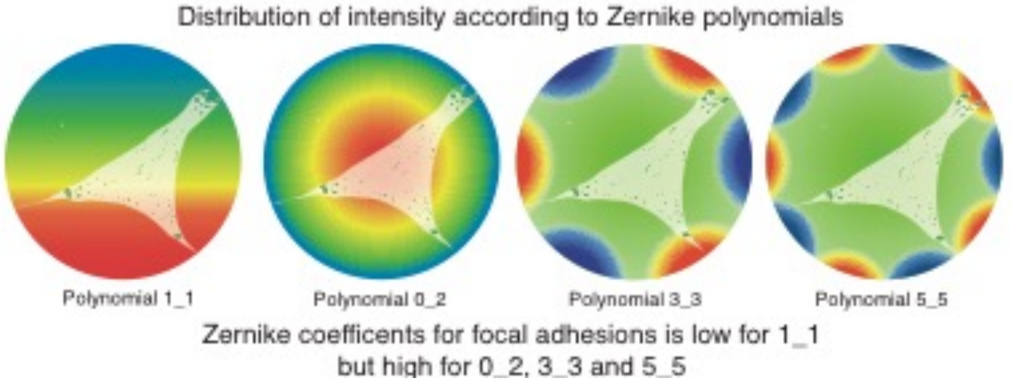<br></td></tr>
<tr><td>Zernike Phase</td><td>Provides information about the orientation of the Zernike magnitude measurement.</td></tr>
<tr><td colspan="3">Haralick texture features<br>The Haralick texture features measure pixel frequency information in an object and this information is summarized in each Haralick measure.</td></tr>
<tr><td>Angular Second Moment</td><td colspan="2">Measures object homogeneity in pixel gray levels. A high value indicates very similar pixels within the vicinity. Based on Haralick's measure, H1</td></tr>
</table>

| | |
|---|---|
| Contrast | Measures the local intensity variations between a reference pixel and its neighbors. A high values indicate large differences in intensity. Based on Haralick's measure, H2 |
| Correlation | Indicates the linearity of the object. A high value is obtained when the object contains linear structures. Based on Haralick's measure, H3 |
| Variance (Sum of Squares) | Measures heterogeneity of pixel gray levels in each object. Based on Haralick's measure, H4 |
| Inverse Difference Moment | Measures local homogeneity. A high value indicates uniform local gray levels. Based on Haralick's measure, H5 |
| Sum Variance | Summation of variance measurements across each object, which gives the distribution of size in grains measured by granularity. Based on Haralick's measure, H7 |
| Sum Entropy | Summation of entropy measurements across each object, which is low for uniform objects. Based on Haralick's measure, H8 |
| Entropy | Measures the degree of randomness in the object. Higher entropy values occur when there is more heterogeneity in neighboring pixels. Based on Haralick's measure, H9 |
| Difference Variance | Difference in variance measurements across each object. Based on Haralick's measure, H10 |
| Difference Entropy | Difference in entropy measurements across each object, which is high for uniform objects. Based on Haralick's measure, H11 |
| Information Measure of Correlation 1 | Ratio of Entropy to inverse difference moment/local homogeneity of the object. Measure of the uncertainty in correlation measurements. Based on Haralick's measure, H12 |
| Information Measure of Correlation 2 | Value related to the square root of entropy. Measure of the uncertainty in correlation measurements. Based on Haralick's measure, H13 |
| Granularity | Measures the grain size. High values indicate the coarseness of the texture. |
| **Non-Haralick textures** | |
| Gabor factor | Measures correlation between bands of intensities. A higher correlation of pattern of intensity results in a higher Gabor feature measurement (e.g. parallel actin stress fibers lead to a higher Gabor factor value) |


[1] L. Kamentsky, T.R. Jones, A. Fraser, M.-A. Bray, D.J. Logan, K.L. Madden, et al., Improved structure, function and compatibility for CellProfiler: modular high-throughput image analysis software, Bioinformatics. 27 (2011) 1179–1180. doi:10.1093/bioinformatics/btr095.

[2] R.M. Kumar, K. Sreekumar, A survey on image feature descriptors, Int J Comput Sci Inf Technol. (2014).

[3] S.L. Vega, A. Dhaliwal, V. Arvind, P.J. Patel, N.R.M. Beijer, J. de Boer, et al., Organizational metrics of interchromatin speckle factor domains: integrative classifier for stem cell adhesion & lineage signaling, Integr. Biol. 7 (2015) 435–446. doi:10.1039/c4ib00281d.
[4] K. Huang, R.F. Murphy, From quantitative microscopy to automated image understanding, J Biomed Opt. 9 (2004) 893–912. doi:10.1117/1.1779233.

| **Table S2. Correlation coefficient of the morphome hierarchically clustered by nanotopography.** | | | | | |
|---|---|---|---|---|---|
| | All topographies | FLAT | SQ | NSQ | HEX |
| All topographies | 1 | | | | |
| FLAT | 0.356 | 1 | | | |
| SQ | 0.389 | 0.140 | 1 | | |
| NSQ | 0.564 | 0.276 | 0.328 | 1 | |
| HEX | 0.458 | 0.194 | 0.229 | 0.451 | 1 |

**Table S3. Mean absolute error (MAE) and coefficient of correlation ($R^2$) of the predicted gene expression values using the morphome.**

| | Test set | |
|---|---|---|
| Gene | MAE | R2 |
| *MYOD1* | 0.10 | 0.10 |
| *MYOG* | 0.10 | 0.10 |
| *MYH7* | 0.10 | 0.10 |
| *RUNX2* | 0.12 | 0.12 |
| *SP7* | 0.17 | 0.17 |
| *BGLAP* | 0.14 | 0.14 |
| *SPP1* | 0.18 | 0.18 |
| *SOX9* | 0.15 | 0.15 |
| *COL2A1* | 0.16 | 0.15 |
| *ACAN* | 0.12 | 0.16 |
| *COL10A* | 0.16 | 0.17 |
| *TGFB1I1* | 0.17 | 0.17 |
| *ELN* | 0.17 | 0.21 |
| *COL3A1* | 0.21 | 0.59 |
| All Genes | 0.11 | 0.72 |

| **Table S4. Sequence of primers used for quantitative measurement of gene expression.** | | | |
|---|---|---|---|
| **Gene** | **Primer sequence** | | **Amplicon size** |
| 18s ribosomal RNA (reference gene) | Fwd | AAGTCCCTGCCCTTTGTACACA | 100 |
| | Rev | GATCCGAGGGCCTCACTAAAC | |
| RUNX2 (Runx2) | Fwd | AAGTGCGGTGCAAACTTTCT | 90 |
| | Rev | TCTCGGTGGCTGCTAGTGA | |
| BGLAP (Osteocalcin) | Fwd | CTGACCTCACAGATGCCAAG | 98 |
| | Rev | GTAGCGCCGGAGTCTGTTC | |
| SPP1 (Osteopontin) | Fwd | TCAGGACAACAACGGAAAGGG | 139 |
| | Rev | GGAACTTGCTTGACTATCGATCAC | |
| SP7 (Osterix) | Fwd | GGTCCAGGCAACACACCTAC | 184 |
| | Rev | GGTAGGGAGCTGGGTTAAGG | |
| SOX9 (Runx2) | Fwd | GGAAGGGAGAGAGAGAGAGAAA | 138 |
| | Rev | CGGGATTTAAGGCTCAAGGT | |
| COL2A1 (Collagen type 2 alpha 1 chain) | Fwd | ACGAAGCGGCTGGCAACCTCA | 73 |
| | Rev | CCCTCGGCCCTCATCTCTACATCA | |
| COL10A1 (Collagen type X alpha 1 chain) | Fwd | TTCTCCTACCACGTGCATGTG | 191 |
| | Rev | AGGCCGTTTGATTCTGCATT | |
| ACAN (Aggrecan) | Fwd | GTGAGGACCTGGTAGTGCGAGTGA | 103 |
| | Rev | GAGCCTGGGCGATAGTGGAATATA | |
| MYOG (Myogenin) | Fwd | GAGACATCCCCCTATTTCTACCA | 106 |
| | Rev | GCTCAGTCCGCTCATAGCC | |
| MYOD1 (Myogenic differentiation 1) | Fwd | CCACTCCGGGACATAGACTTG | 109 |
| | Rev | AAAAGCGCAGGTCTGGTGAG | |
| MYH7 (Myosin heavy chain 7) | Fwd | CTCAAGCTGCTCAGCAATCTATTT | 153 |
| | Rev | GGAGCGCAAGTTTGTCATAAGT | |
| COL3A1 (Collagen type III alpha chain 1) | Fwd | CTGTAACATGGAAACTGGGGAAA | 144 |
| | Rev | CCATAGCTGAACTGAAAACCACC | |
| | Fwd | ATGTCACGGTTAGGGGCTC | 83 |

| | | | |
|---|---|---|---|
| TGFB1I1 (Transforming growth factor beta 1 induced transcript 1) | Rev | GGCTTGCATACTGTGCTGTATAG | |
| ELN (Elastin) | Fwd | TGTCCCACTGGGTTATCCCAT | 92 |
| | Rev | CAGCTACTCCATAGGGCAATTTC | |
