## Supplemental Data 2 for "Predicting gene expression using morphological cell responses to nanotopography"

### Supplementary methods

**Conventional cell measurements**

Images of cells stained against the chromatin, actin, focal adhesion kinase (FAK) and FAK phosphorylated at the Tyrosine 397 site (pFAK) were used to measure cell area, nucleus area, total pFAK/FAK integrated intensity ratio, actin integrated intensity and individual focal adhesion area. The nuclei was segmented using the image of chromatin, the cell body was segmented using the image of actin, while individual focal adhesions were segmented using the image of pFAK.

**Logistic regression**

Logistic regression was used as a supervised classification algorithm to separate the morphome according to cell type or nanotopography. Logistic regression was performed using a 10-fold cross validation method[1-3], where the entire data is separated into 10 subsets. At each iteration, the logistic regressor is trained using 9 out of 10 subsets and tested using the held-out subset. The procedure is repeated across all subsets. Accuracy of the logistic regressor is presented as the mean ± standard deviation measured from the test set across 10-folds. A receiver operating characteristic (ROC) curve using a one class-vs-all analysis was created from the logistic regressor with the highest level of accuracy measured using the test set. Logistic regression and ROC was created using the *nnet*[4] and *caret*[5] package, respectively, on R.

**Mineralization study**

After 28 days, calcium phosphate deposition was studied using Alizarin red and Von Kossa histological stains. Cells were fixed with 4% paraformaldehyde and stained against 4% Alizarin red (pH 4.2, Sigma Aldrich), or von Kossa (Merck Millipore) and NuclearFast Red (Sigma Aldrich). Samples were imaged at 10X magnification using a color CCD camera.
