## Supplemental Data 3 for "Predicting gene expression using morphological cell responses to nanotopography"

### Supplementary figures

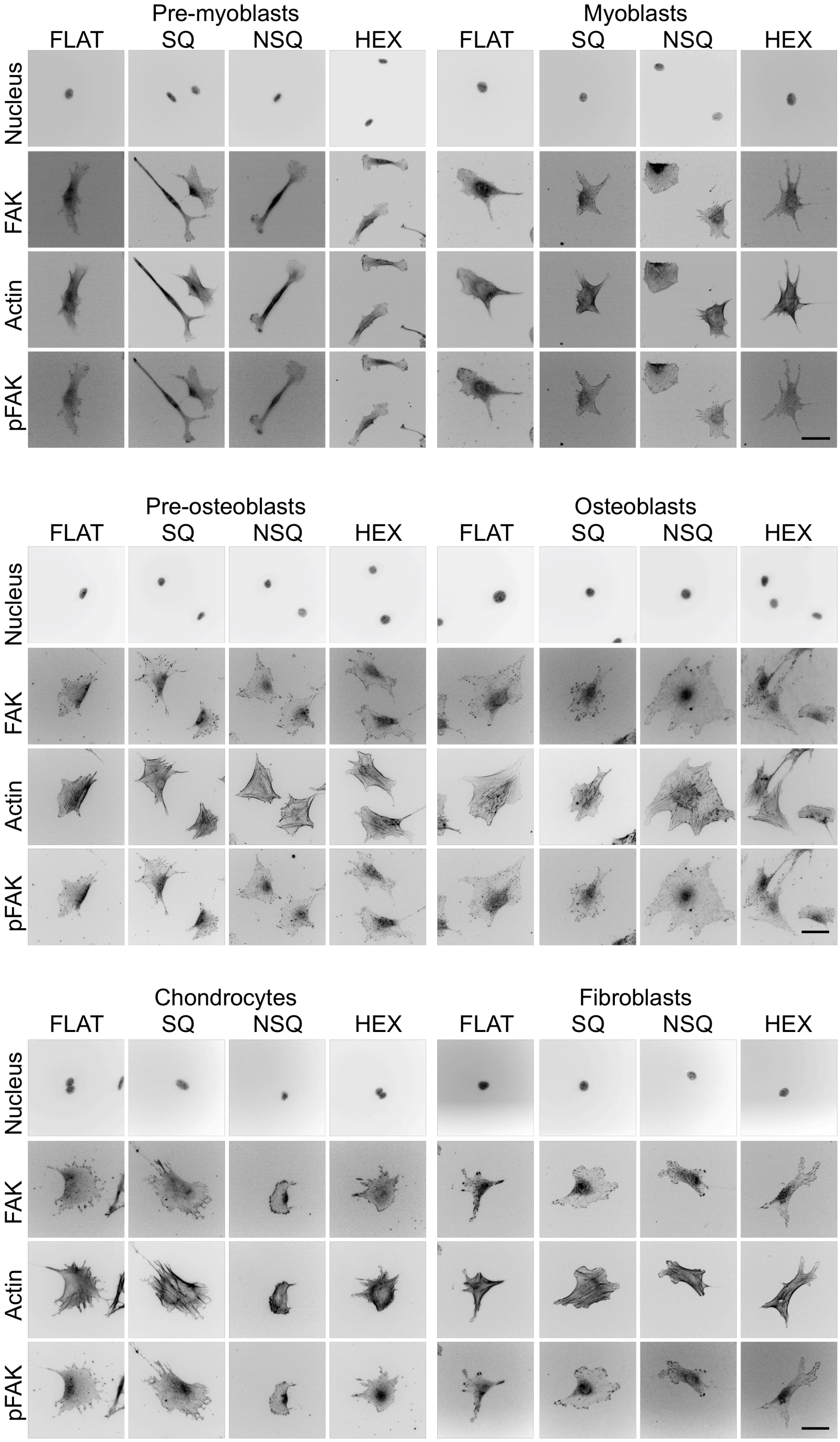
Pre-myoblasts
Myoblasts
FLAT
SQ
NSQ
HEX
FLAT
SQ
NSQ
HEX
Nucleus
FAK
Actin
pFAK
Pre-osteoblasts
Osteoblasts
FLAT
SQ
NSQ
HEX
FLAT
SQ
NSQ
HEX
Nucleus
FAK
Actin
pFAK
Chondrocytes
Fibroblasts
FLAT
SQ
NSQ
HEX
FLAT
SQ
NSQ
HEX
Nucleus
FAK
Actin
pFAK

Figure S1. Representative images of musculoskeletal cell types cultured on four different nanotopographies. Cells were stained using antibodies against the nucleus, actin cytoskeleton, focal adhesion kinase (FAK) and phosphorylated (Y397) focal adhesion kinase (pFAK) and imaged after 2 days in culture on the nanotopographies: FLAT, SQ, NSQ, and HEX. Scale bar = 50 μm.

Cells manifested qualitative differences after 2 days. Pre-myoblasts and myoblasts were highly elongated on FLAT but were more irregularly shaped with highly discrete and large focal adhesions on SQ. Pre-osteoblasts and osteoblasts had very prominent actin stress fibers and large adhesions regardless of topography. Chondrocytes generally had low pFAK expression, weak stress fiber formation, and stellate and irregular shapes, particularly on SQ and NSQ. Focal adhesions were largest in osteoblastic cells on NSQ and SQ. Fibroblasts had very diffuse focal adhesions and were small and elongated on FLAT and SQ, while those on NSQ and HEX were larger and with more aligned actin stress fibers.

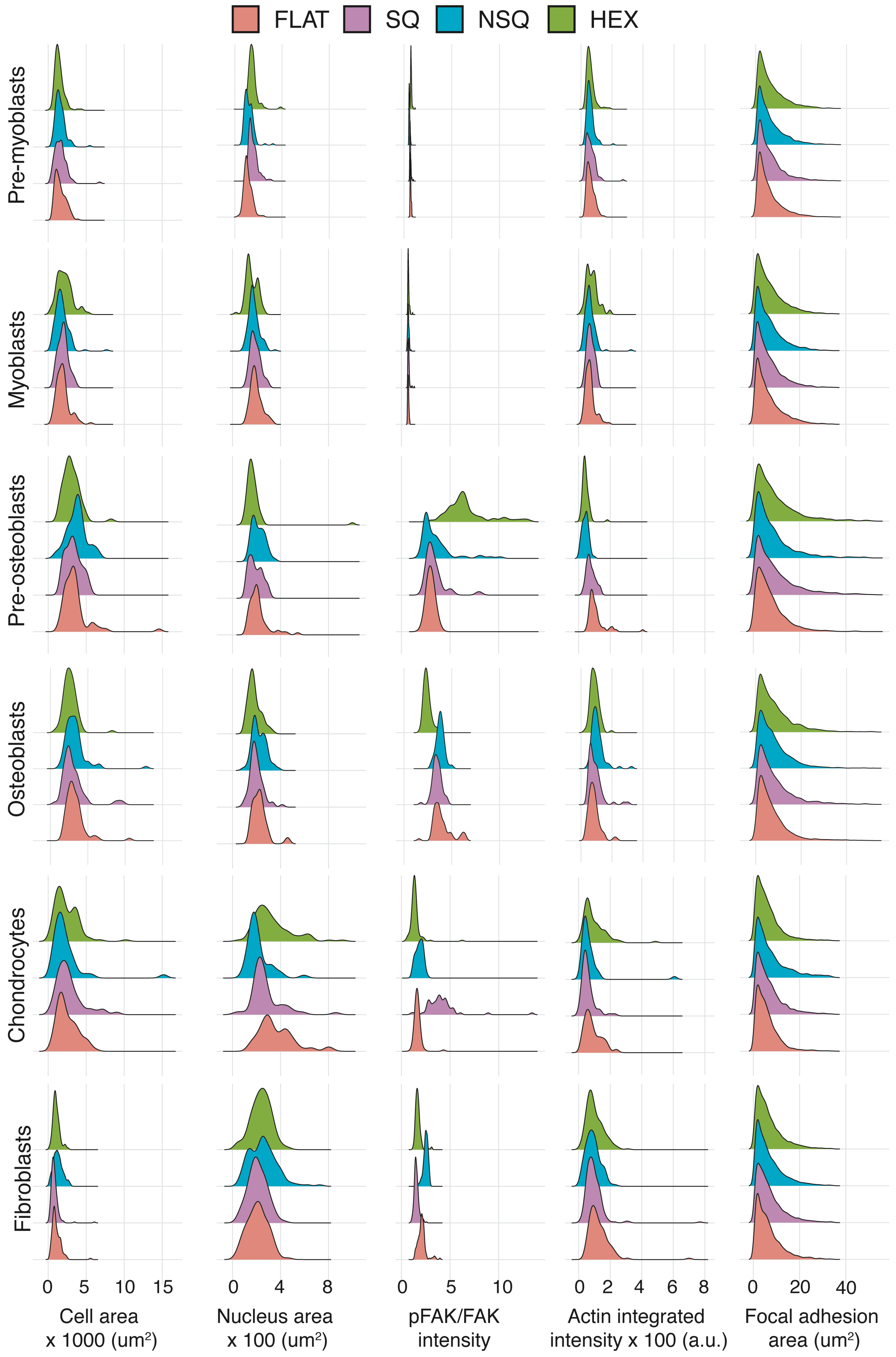
FLAT
SQ
NSQ
HEX
Pre-myoblasts
Myoblasts
Pre-osteoblasts
Osteoblasts
Chondrocytes
Fibroblasts
0 5 10 15
Cell area x 1000 (um²)
0 4 8
Nucleus area x 100 (um²)
0 5 10
pFAK/FAK intensity
0 2 4 6 8
Actin integrated intensity x 100 (a.u.)
0 20 40
Focal adhesion area (um²)

Figure S2. Conventional measurements of cells and focal adhesions reveal cell type specific changes induced by nanotopography. One-way ANOVA showed statistically significant differences across all cell type and topography combinations for cell area, nucleus area, pFAK/FAK integrated intensity ratio, actin integrated intensity and focal adhesion area. P values adjusted for multiple comparisons with Tukey's post-hoc test is listed in Supplementary data (Statistical analysis – conventional measurements). NSQ significantly increased cell area relative to FLAT. Changes in pFAK/FAK intensity ratios showed cell type dependence: chondrocytes cultured on SQ, pre-osteoblasts cultured on HEX, and fibroblasts on HEX showed the highest pFAK activation compared to the FLAT control. Generally, both pre-myoblasts and myoblasts showed low pFAK activation compared to all cell types. Actin intensity changed most markedly in chondrocytes cultured on HEX, showing higher variance and higher intensity values compared with the FLAT control. Pre-osteoblasts and osteoblasts on NSQ and HEX developed focal adhesions with the largest area compared to all cell types.m a

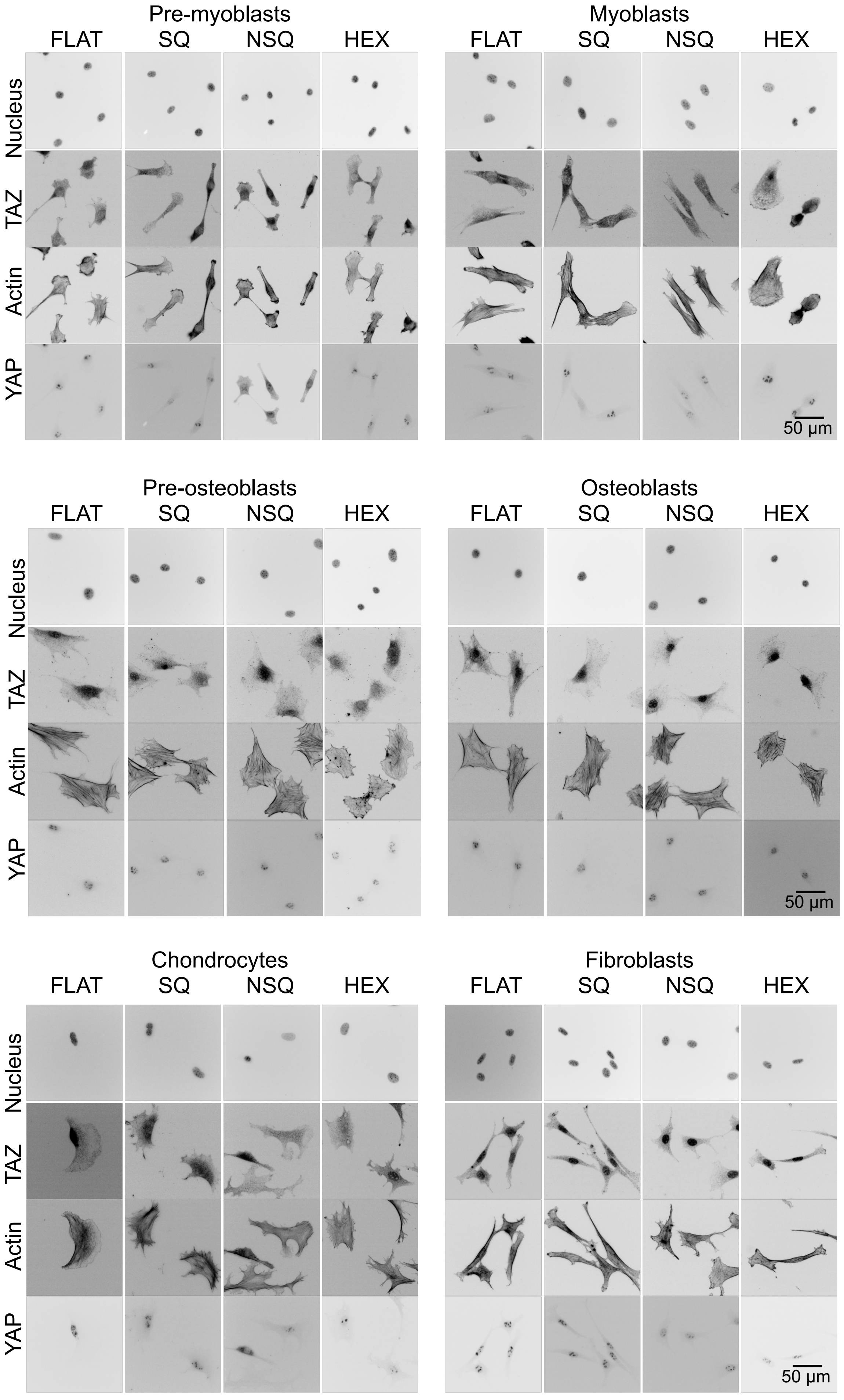
Pre-myoblasts
FLAT
SQ
NSQ
HEX
Myoblasts
FLAT
SQ
NSQ
HEX
Nucleus
TAZ
Actin
YAP
50 µm
Pre-osteoblasts
FLAT
SQ
NSQ
HEX
Osteoblasts
FLAT
SQ
NSQ
HEX
Nucleus
TAZ
Actin
YAP
50 µm
Chondrocytes
FLAT
SQ
NSQ
HEX
Fibroblasts
FLAT
SQ
NSQ
HEX
Nucleus
TAZ
Actin
YAP
50 µm

Figure S3. Representative images of cellular YAP/TAZ expression and localization in response to nanotopography. Cells were stained using antibodies against the nucleus, actin cytoskeleton, and the mechanosensors YAP and TAZ then imaged after 2 days in culture on the nanotopographies. Scale bar = 50 µm.

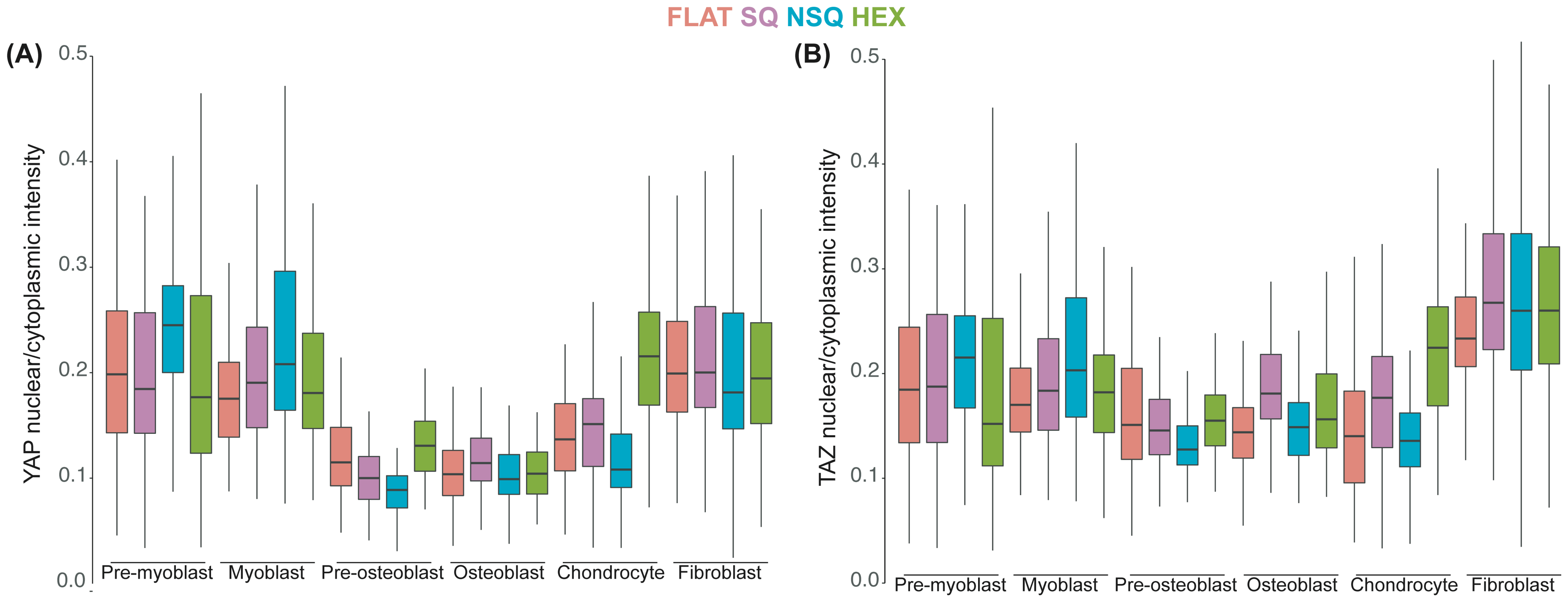


Figure S4. Quantification of YAP/TAZ nuclear localization changed by nanotopography. The ratio of integrated intensity in the nucleus normalized to total intensity in the cell of (A) YAP or (B) TAZ are shown here. Changes in the nuclear/cytoplasmic intensities of YAP or TAZ indicate changes in mechanosignaling induced by nanotopographical cues. The effects of nanotopography on YAP or TAZ nuclear translocation vary according to the cell type. Relative to its FLAT controls, NSQ induced higher whereas SQ showed lower nuclear translocation of YAP and TAZ on myoblastic and osteoblastic cells, respectively.

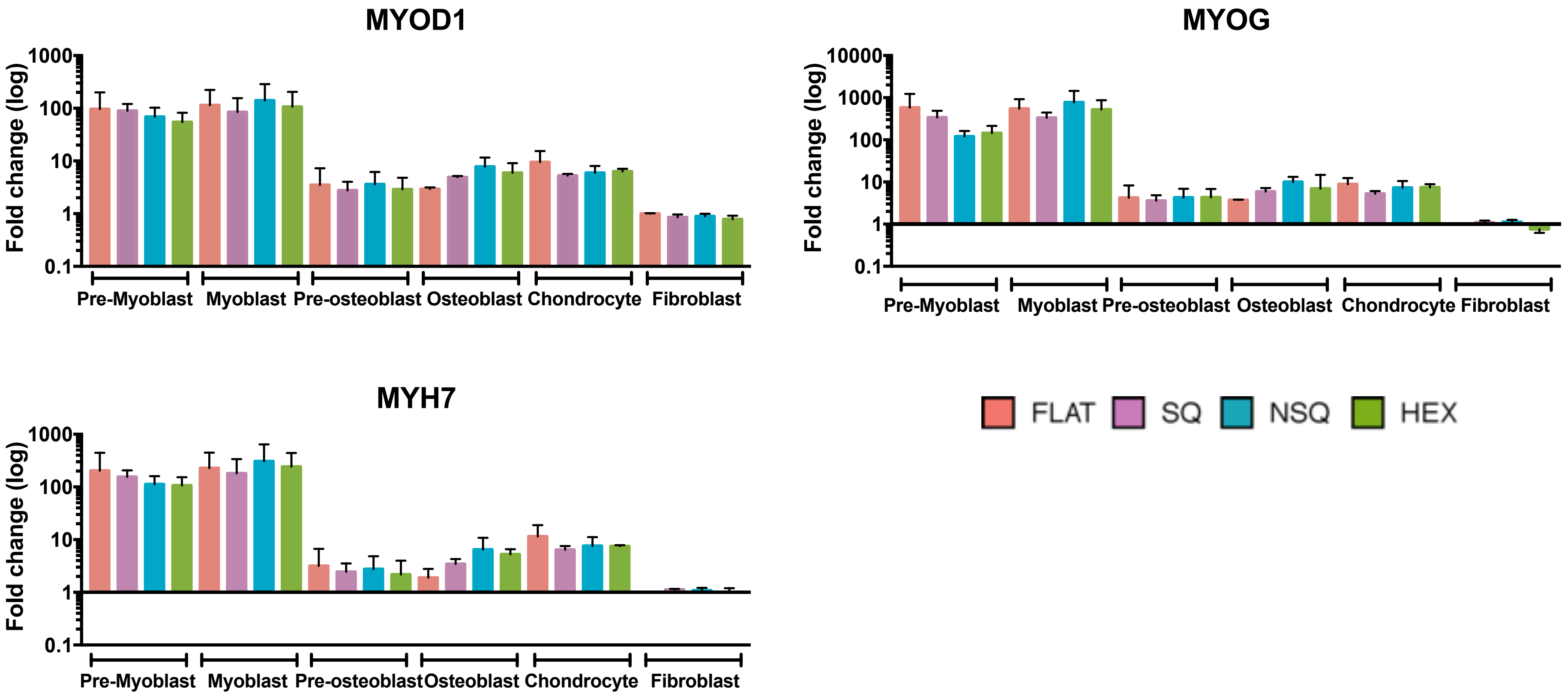
MYOD1
MYOG
MYH7
Fold change (log)
1000
100
10
1
0.1
10000
Pre-Myoblast
Myoblast
Pre-osteoblast
Osteoblast
Chondrocyte
Fibroblast
FLAT
SQ
NSQ
HEX

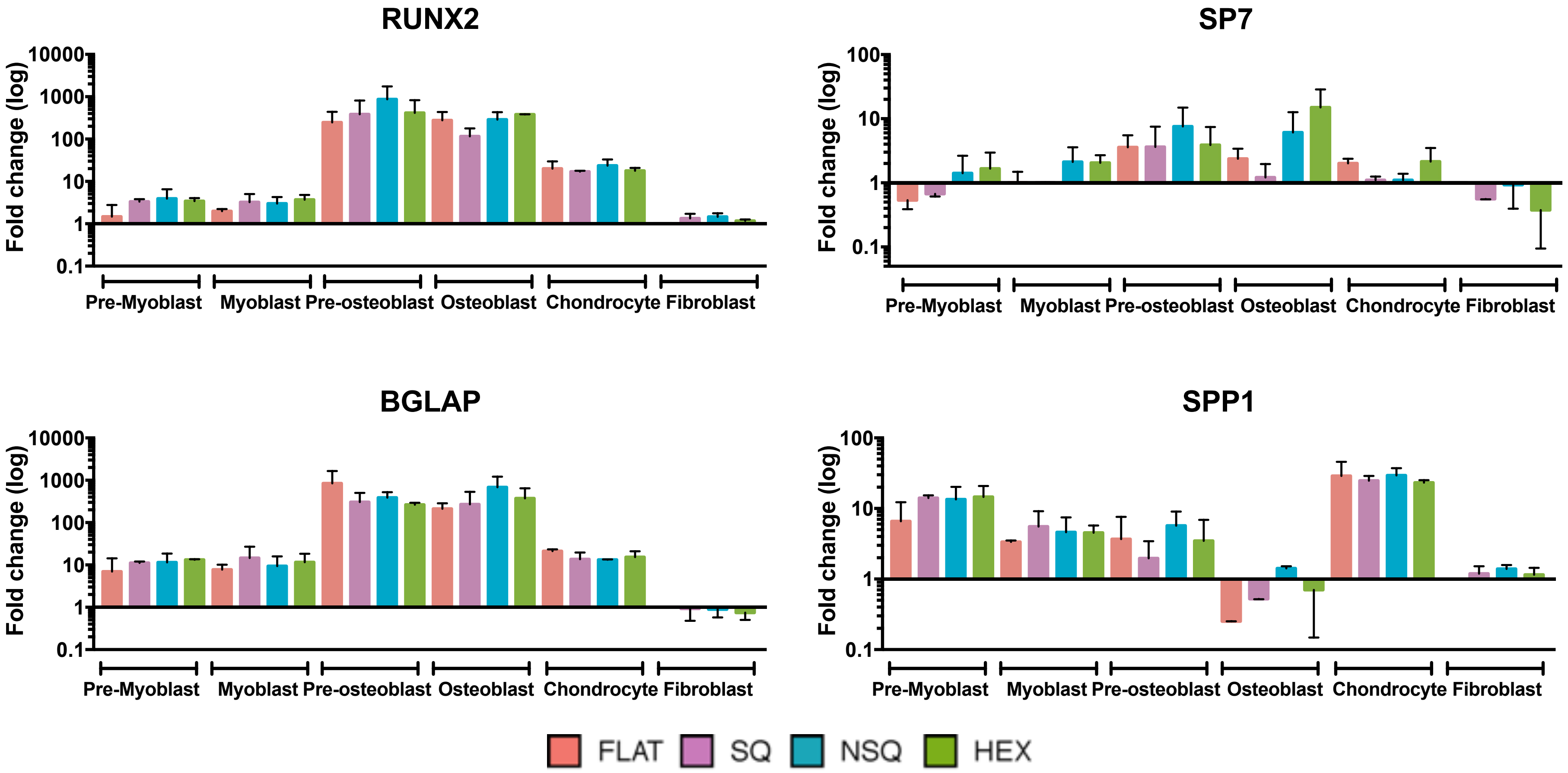
RUNX2
SP7
BGLAP
SPP1
Fold change (log)
10000
1000
100
10
1
0.1
Pre-Myoblast
Myoblast
Pre-osteoblast
Osteoblast
Chondrocyte
Fibroblast
FLAT
SQ
NSQ
HEX

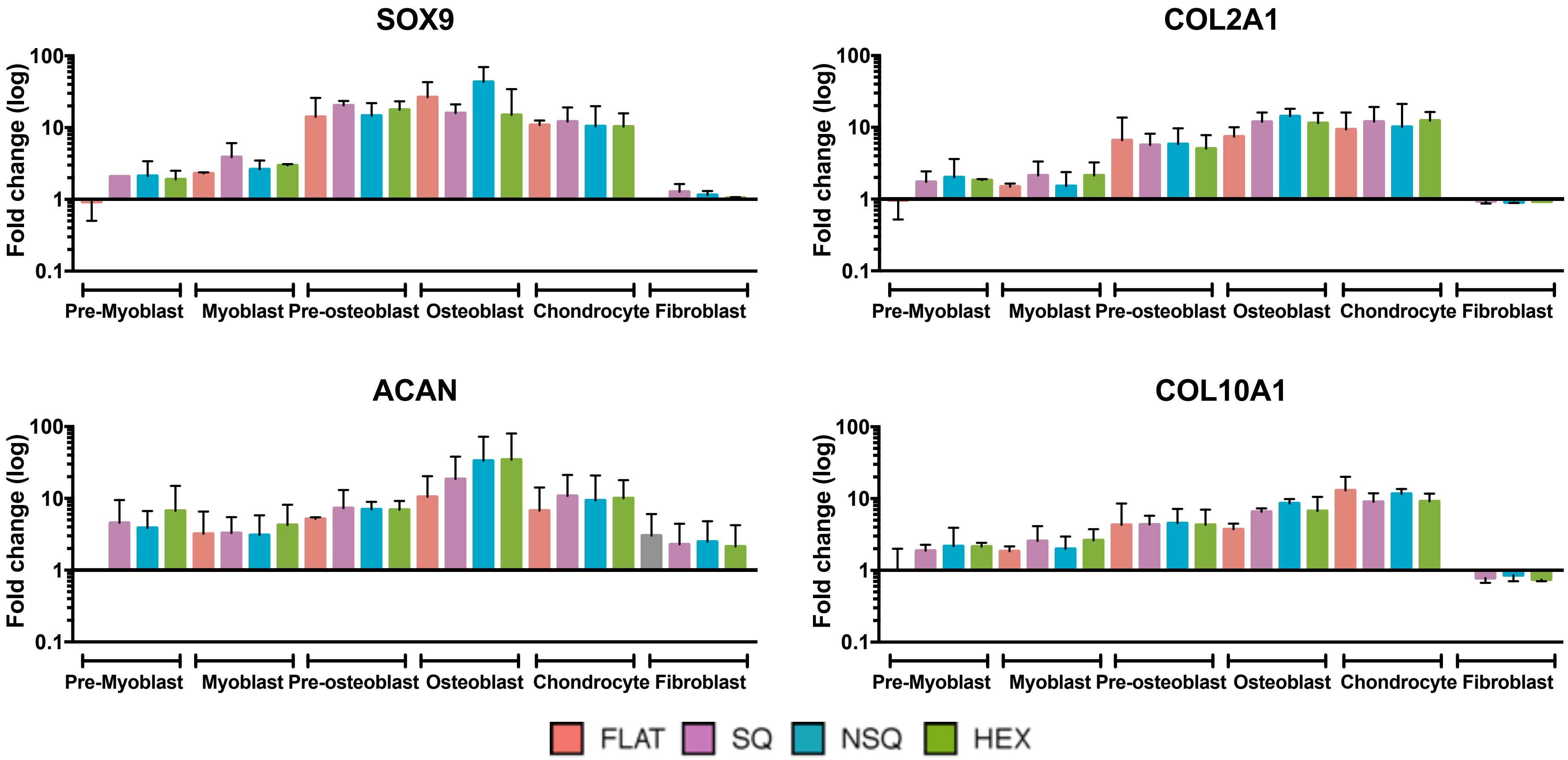
SOX9
COL2A1
ACAN
COL10A1
Fold change (log)
100
10
1
0.1
Pre-Myoblast
Myoblast
Pre-osteoblast
Osteoblast
Chondrocyte
Fibroblast
FLAT
SQ
NSQ
HEX

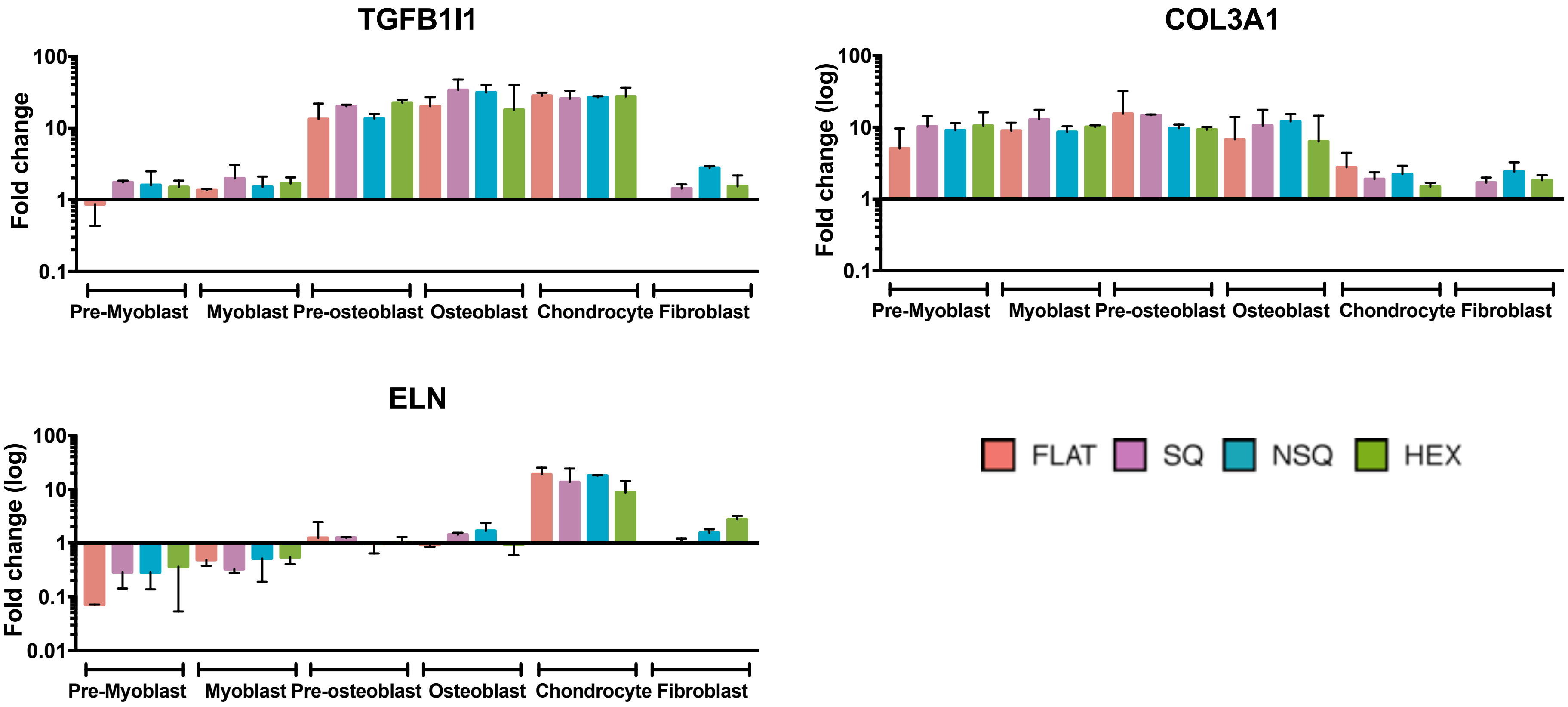


Figure S5. Measurement of musculoskeletal marker expression on all combinations of cell type and topography. Quantitative polymerase chain reaction (qPCR) was used to determine the changes in gene expression across 14 different musculoskeletal genes. Fold change in expression of each gene expression were calculated with respect to fibroblasts on Flat controls (n=6).

(A)

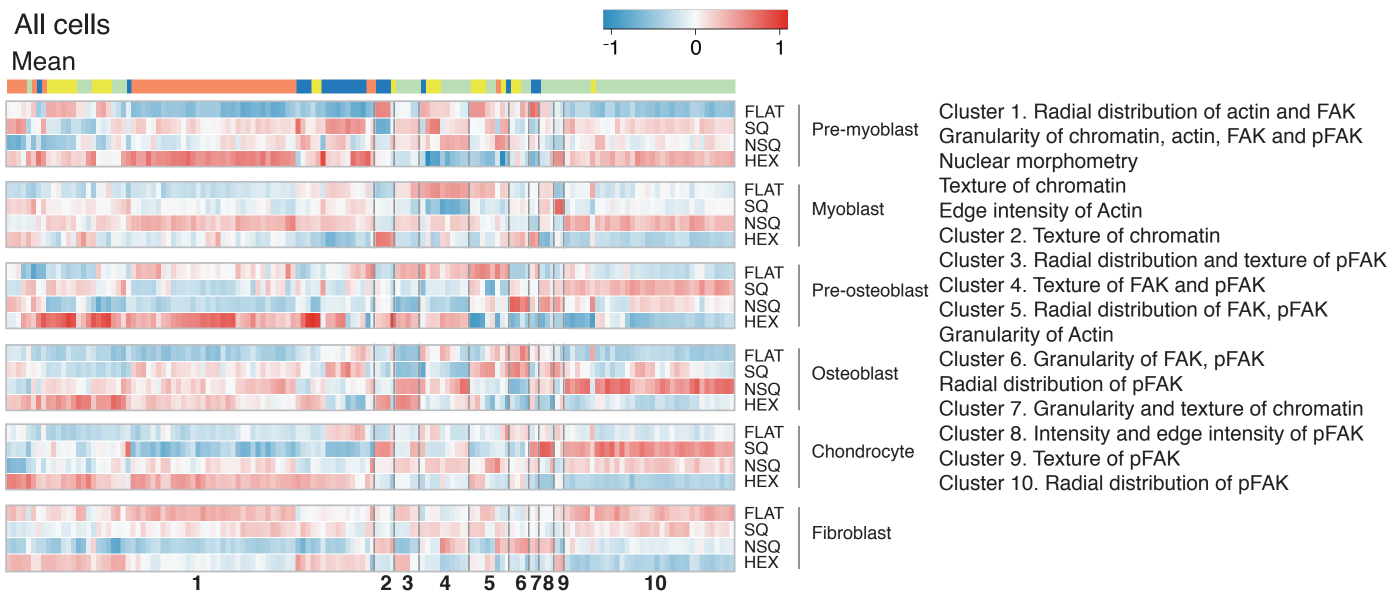


(B)

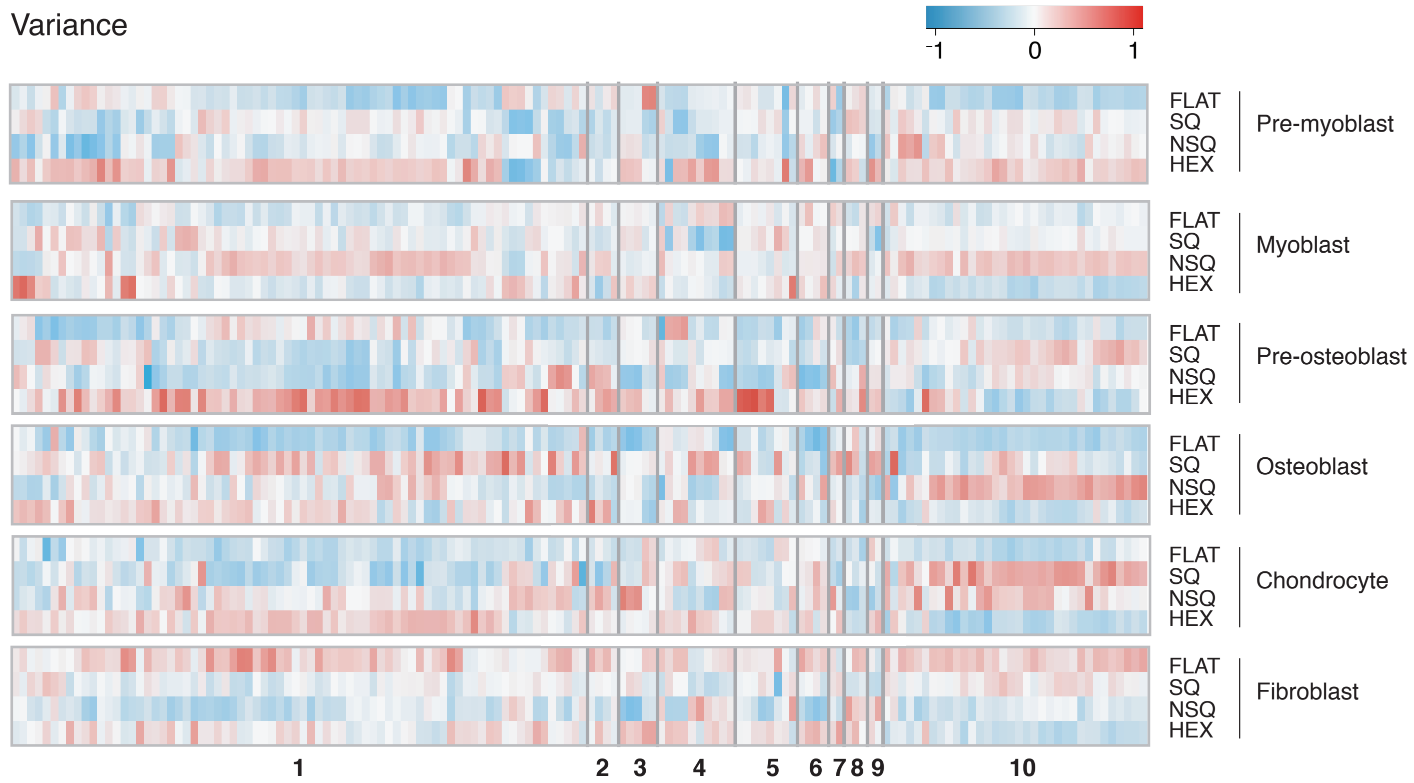


Figure S6. Hierarchical clustering of the (A) mean measurements and (B) variance in the morphome across nanotopographies. For visualization, each morphome feature was normalized to have a mean value of 0 and standard deviation of 1 across all datapoints. Features included were significantly varied across cell types ($p < 0.05$ using one-way ANOVA).

(B)

| Actual | | | | | | |
|---|---|---|---|---|---|---|
| | **Pre-myoblast** | **Myoblast** | **Pre-osteoblast** | **Osteoblast** | **Chondrocyte** | **Fibroblast** |
| **Pre-myoblast** | 99 ± 1.04 | 0.11 ± 0.34 | 1.09 ± 1.04 | 0.14 ± 0.43 | 0.48 ± 1.1 | 0.09 ± 0.28 |
| **Myoblast** | 0.11 ± 0.36 | 99 ± 0.79 | 0.31 ± 0.66 | 0.14 ± 0.44 | 0.16 ± 0.52 | 0.35 ± 0.45 |
| **Pre-osteoblast** | 0.57 ± 0.80 | 0.32 ± 0.52 | 98 ± 1.68 | 0.14 ± 0.44 | 0.16 ± 0.51 | 0.26 ± 0.59 |
| **Osteoblast** | 0.11 ± 0.36 | 0.22 ± 0.45 | 0.16 ± 0.49 | 99 ± 1.52 | 0.16 ± 0.51 | 0.09 ± 0.28 |
| **Chondrocyte** | 0.11 ± 0.36 | 0.11 ± 0.34 | 0 ± 0 | 0.41 ± 0.66 | 99 ± 1.90 | 0.18 ± 0.37 |
| **Fibroblast** | 0.46 ± 0.59 | 0.43 ± 0.56 | 0.78 ± 0.82 | 0.41 ± 0.94 | 0.48 ± 0.78 | 99 ± 0.97 |

Predicted

(B)

Figure S7. Classification of different cell types using the morphome. A supervised machine learning method (logistic regression) was trained to distinguish 6 different cell types. (A) Confusion matrix showing high classification accuracies of the logistic regression model for classifying 6 different cell types. Data are presented as mean ± standard deviation from a 10-fold cross validation. (B) The receiver operating characteristic (ROC) curve showing performance of the logistic regression model for classifying the data into 6 different cell types. The numbers denote the area under the ROC curve, indicating robustness of the model.

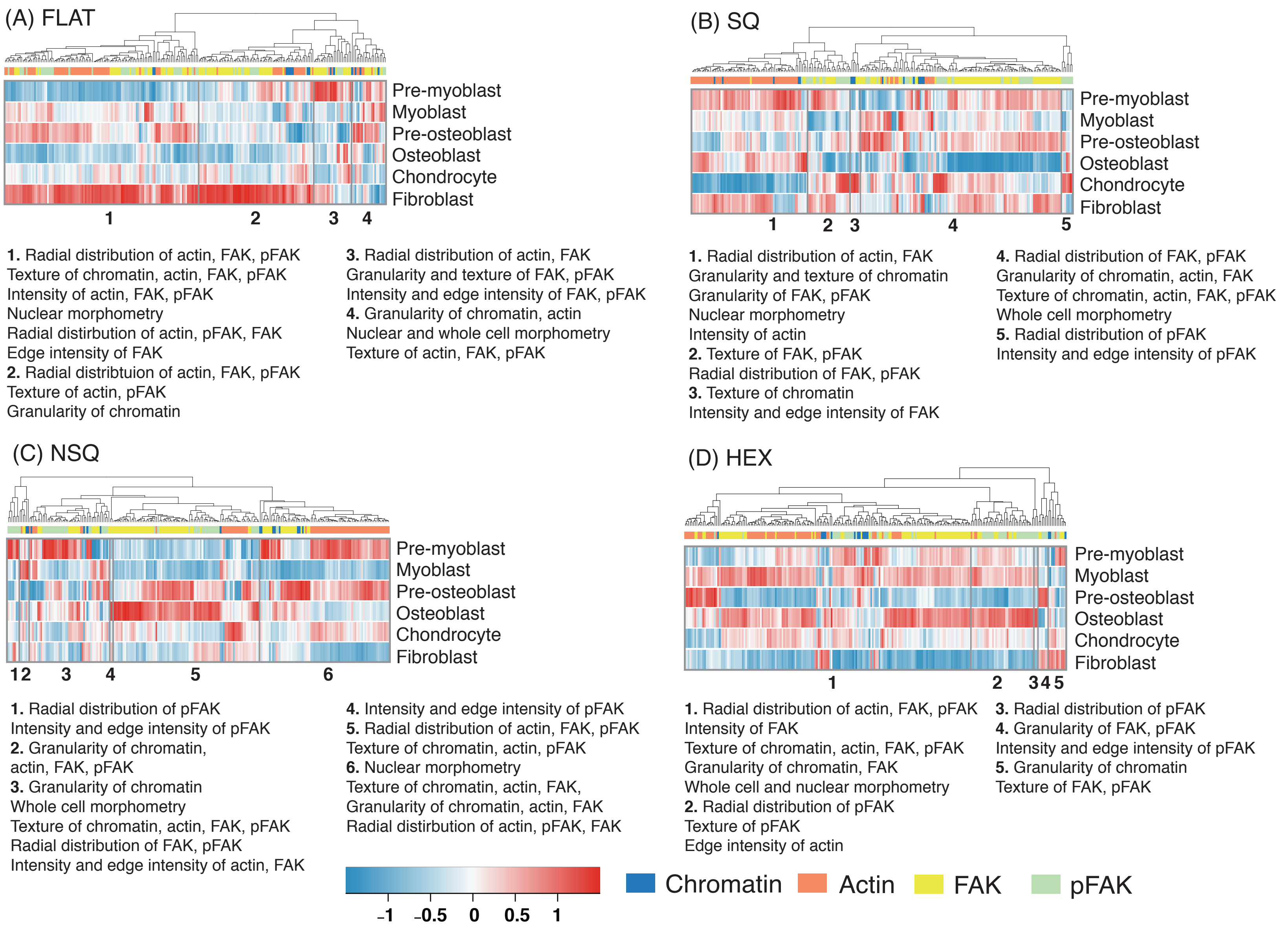
(A) FLAT
Pre-myoblast
Myoblast
Pre-osteoblast
Osteoblast
Chondrocyte
Fibroblast
1 2 3 4
1. Radial distribution of actin, FAK, pFAK
Texture of chromatin, actin, FAK, pFAK
Intensity of actin, FAK, pFAK
Nuclear morphometry
Radial distirbution of actin, pFAK, FAK
Edge intensity of FAK
2. Radial distribtuion of actin, FAK, pFAK
Texture of actin, pFAK
Granularity of chromatin
3. Radial distribution of actin, FAK
Granularity and texture of FAK, pFAK
Intensity and edge intensity of FAK, pFAK
4. Granularity of chromatin, actin
Nuclear and whole cell morphometry
Texture of actin, FAK, pFAK
(B) SQ
Pre-myoblast
Myoblast
Pre-osteoblast
Osteoblast
Chondrocyte
Fibroblast
1 2 3 4 5
1. Radial distribution of actin, FAK
Granularity and texture of chromatin
Granularity of FAK, pFAK
Nuclear morphometry
Intensity of actin
2. Texture of FAK, pFAK
Radial distribution of FAK, pFAK
3. Texture of chromatin
Intensity and edge intensity of FAK
4. Radial distribution of FAK, pFAK
Granularity of chromatin, actin, FAK
Texture of chromatin, actin, FAK, pFAK
Whole cell morphometry
5. Radial distribution of pFAK
Intensity and edge intensity of pFAK
(C) NSQ
Pre-myoblast
Myoblast
Pre-osteoblast
Osteoblast
Chondrocyte
Fibroblast
1 2 3 4 5 6
1. Radial distribution of pFAK
Intensity and edge intensity of pFAK
2. Granularity of chromatin,
actin, FAK, pFAK
3. Granularity of chromatin
Whole cell morphometry
Texture of chromatin, actin, FAK, pFAK
Radial distribution of FAK, pFAK
Intensity and edge intensity of actin, FAK
4. Intensity and edge intensity of pFAK
5. Radial distribution of actin, FAK, pFAK
Texture of chromatin, actin, pFAK
6. Nuclear morphometry
Texture of chromatin, actin, FAK,
Granularity of chromatin, actin, FAK
Radial distirbution of actin, pFAK, FAK
(D) HEX
Pre-myoblast
Myoblast
Pre-osteoblast
Osteoblast
Chondrocyte
Fibroblast
1 2 3 4 5
1. Radial distribution of actin, FAK, pFAK
Intensity of FAK
Texture of chromatin, actin, FAK, pFAK
Granularity of chromatin, FAK
Whole cell and nuclear morphometry
2. Radial distribution of pFAK
Texture of pFAK
Edge intensity of actin
3. Radial distribution of pFAK
4. Granularity of FAK, pFAK
Intensity and edge intensity of pFAK
5. Granularity of chromatin
Texture of FAK, pFAK
–1 –0.5 0 0.5 1
Chromatin
Actin
FAK
pFAK

(caption for figure in previous page) Figure S8. Morphome separated by nanotopographies clustered characteristically across each cell type. (A-D) Hierarchical clustering of the morphome separated by nanotopography. Cell type specific morphome features are made up of 15 chromatin, 73 actin, 88 FAK and 56 pFAK features. Each morphome feature was mean centered and normalized to standard deviation per cell type to result in mean = 0 and standard deviation = 1. The total number of cells analysed per nanotopography type are: (A) n=1251 for FLAT; (B) n=1215 for SQ; (C) n=1293 for NSQ; (D) n=1180 for HEX obtained across 2 biological replicates. The colour and intensity of each tile represents the average value of the feature for the specified cell type and nanotopography.

On FLAT, pre-myoblasts and osteoblasts showed predominantly low average values for cell-type specific features except for morphometric and texture measurements (cluster 4). This was reversed in fibroblasts, with particularly high average values of FAK and actin radial distribution. Pre-myoblasts showed high average values for actin radial distribution and intensity, nuclear morphometry, and chromatin textures on SQ (cluster 1). Chondrocytes showed the opposite profile for the same features. Average values for FAK measurements were high on pre-myoblasts and pre-osteoblasts and low on osteoblasts (cluster 4). On NSQ, pre-osteoblasts and osteoblasts showed markedly high average values of FAK radial distribution (cluster 5), relative to other cell types while pre-myoblasts and myoblasts cells showed low average values of FAK radial distribution. Pre-myoblasts and myoblasts cells showed the opposite for the same features. While pre-osteoblasts and myoblasts demonstrated high average values of actin radial distribution and textures (cluster 6), osteoblasts and myoblasts showed low average values for the same features. Cell type-specific features were mostly high on pre-myoblasts, myoblasts and osteoblasts on HEX. On the other hand, fibroblasts and pre-osteoblasts showed low average values for the cell-type specific features. Surprisingly, chondrocytes exhibited moderate values (close to the mean) of the cell type-specific features. The cell type-specific hierarchical clusters show low correlation (Table S2).

(B)

| Predicted \ Actual | FLAT | SQ | NSQ | HEX |
|---|---|---|---|---|
| FLAT | 65 ± 5.3 | 17 ± 3.3 | 13 ± 3.2 | 6.5 ± 1.9 |
| SQ | 19 ± 3.4 | 58 ± 4.8 | 18 ± 2.4 | 3.2 ± 1.2 |
| NSQ | 9.7 ± 2.0 | 20 ± 3.1 | 61 ± 3.3 | 3.8 ± 1.4 |
| HEX | 5.9 ± 2.0 | 4.4 ± 1.8 | 7.6 ± 1.8 | 87 ± 2.6 |

(B)

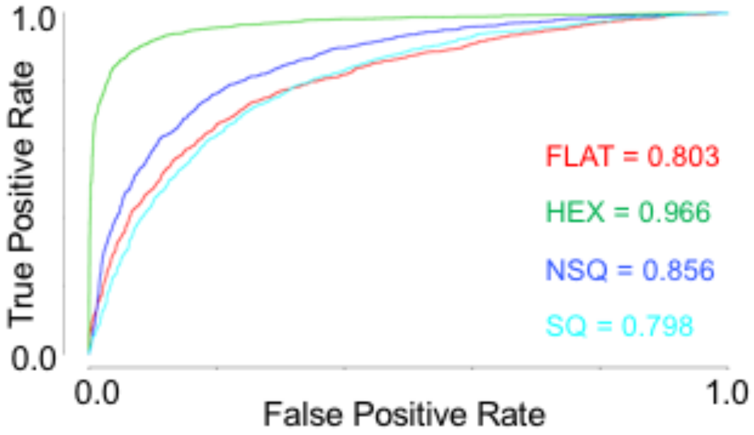


Figure S9. Classification of different nanotopographies using the morphome. A supervised machine learning method (logistic regression) was trained to distinguish 4 different nanotopographies. (A) Confusion matrix showing classification accuracies of the logistic regression model for classifying different nanotopographies. Data are presented as mean ± standard deviation from a 10-fold cross validation. (B) The ROC curve showing performance of the logistic regression model for classifying the data into 6 different nanotopographies. The numbers denote the area under the ROC curve, indicating robustness of the model.

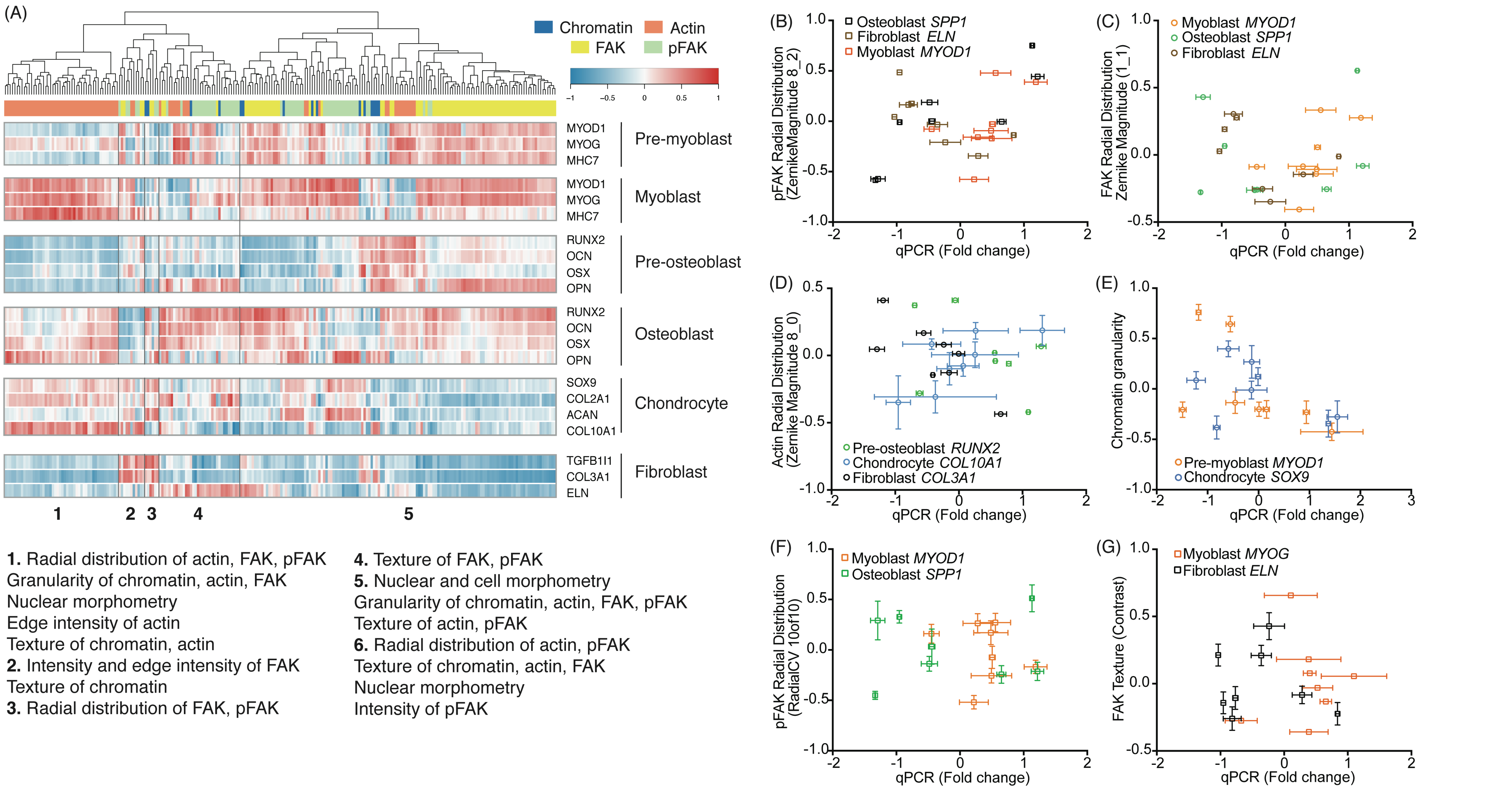
(A)
Chromatin
Actin
FAK
pFAK
MYOD1
MYOG
MHC7
Pre-myoblast
MYOD1
MYOG
MHC7
Myoblast
RUNX2
OCN
OSX
OPN
Pre-osteoblast
RUNX2
OCN
OSX
OPN
Osteoblast
SOX9
COL2A1
ACAN
COL10A1
Chondrocyte
TGFB1I1
COL3A1
ELN
Fibroblast
1 2 3 4 5
1. Radial distribution of actin, FAK, pFAK
Granularity of chromatin, actin, FAK
Nuclear morphometry
Edge intensity of actin
Texture of chromatin, actin
2. Intensity and edge intensity of FAK
Texture of chromatin
3. Radial distribution of FAK, pFAK
4. Texture of FAK, pFAK
5. Nuclear and cell morphometry
Granularity of chromatin, actin, FAK, pFAK
Texture of actin, pFAK
6. Radial distribution of actin, pFAK
Texture of chromatin, actin, FAK
Nuclear morphometry
Intensity of pFAK
(B)
Osteoblast SPP1
Fibroblast ELN
Myoblast MYOD1
pFAK Radial Distribution (ZernikeMagnitude 8_2)
qPCR (Fold change)
(C)
Myoblast MYOD1
Osteoblast SPP1
Fibroblast ELN
FAK Radial Distribution Zernike Magnitude (1_1)
qPCR (Fold change)
(D)
Pre-osteoblast RUNX2
Chondrocyte COL10A1
Fibroblast COL3A1
Actin Radial Distribution (Zernike Magnitude 8_0)
qPCR (Fold change)
(E)
Pre-myoblast MYOD1
Chondrocyte SOX9
Chromatin granularity
qPCR (Fold change)
(F)
Myoblast MYOD1
Osteoblast SPP1
pFAK Radial Distribution (RadialCV 10of10)
qPCR (Fold change)
(G)
Myoblast MYOG
Fibroblast ELN
FAK Texture (Contrast)
qPCR (Fold change)

Figure S10. Spearman correlation between the changes in morphome and changes in gene expression resulting from nanotopography. Spearman correlation coefficients were used to measure the similarities in changes between morphome and gene expression across nanotopographies. (A) Hierarchical clustering of the correlation coefficients for each morphome and gene expression pair. The colour and intensity of each tile represents the correlation coefficient magnitude and direction of association between the morphome feature and gene expression. Each feature was mean centered and normalized to standard deviation. (B-G) Scatterplots showing changes in qPCR against individual morphome features. Spearman rank correlation was performed using KNIME (v2.0).

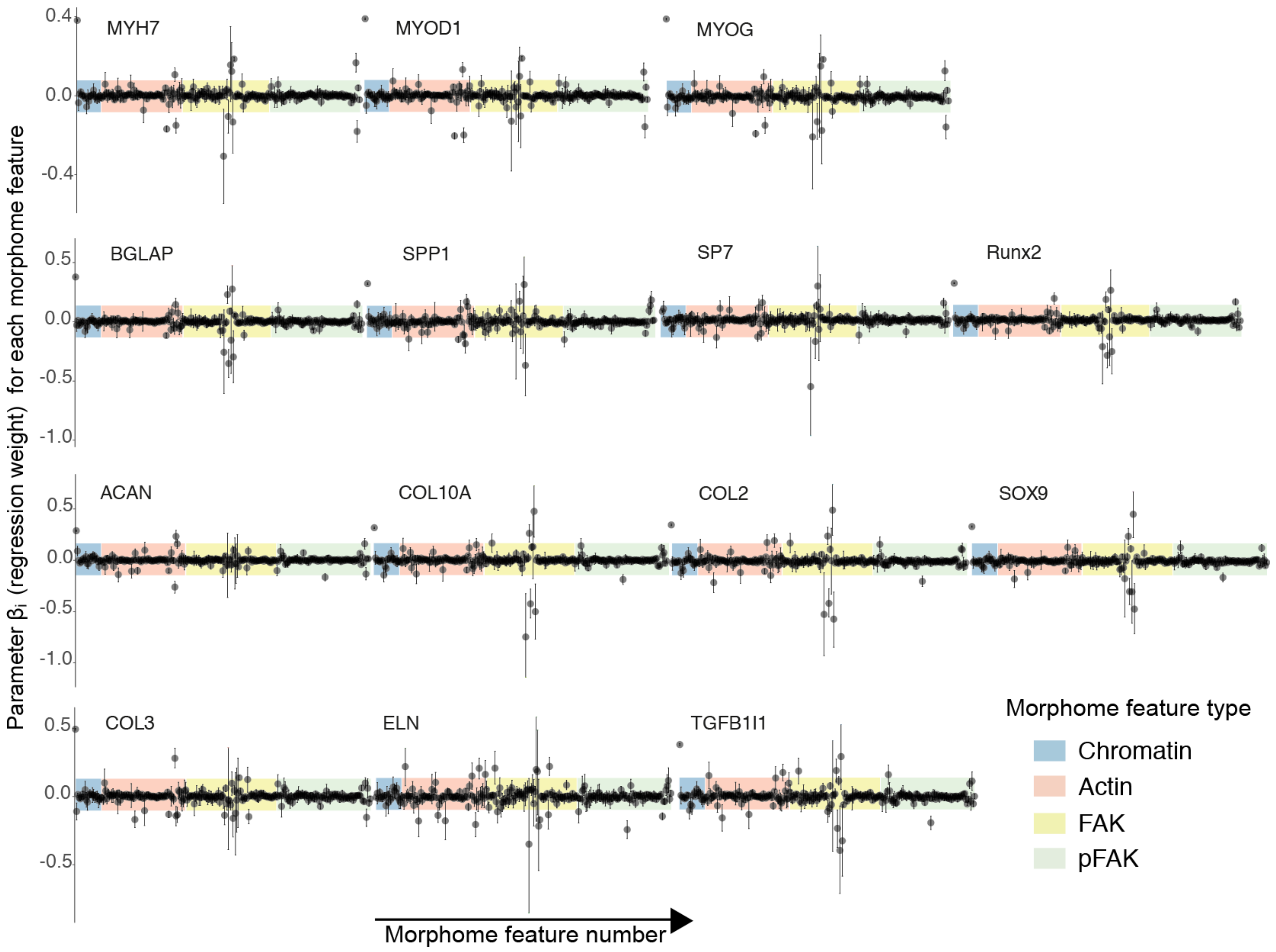

MYH7
MYOD1
MYOG
BGLAP
SPP1
SP7
Runx2
ACAN
COL10A
COL2
SOX9
COL3
ELN
TGFB1I1
0.4
0.0
-0.4
0.5
0.0
-0.5
-1.0
0.5
0.0
-0.5
-1.0
0.5
0.0
-0.5
Parameter βi (regression weight) for each morphome feature
Morphome feature number
Morphome feature type
Chromatin
Actin
FAK
pFAK

Figure S11. Contribution of each morphome feature to the Bayesian linear regression model. The parameters β from the linear regression models (i.e. regression weight) weights the contribution of each morphome feature in the Bayesian linear regression model for predicting gene expression. Higher magnitudes of the parameter β indicate larger contribution of the morphome feature in predicting the expression of the specified gene. Data shown are mean (dots) ± high density interval (bars) of β for each morphome feature and gene. The morphome features were color coded to denote chromatin (blue), actin (orange), FAK (yellow) and pFAK (green) measurements.

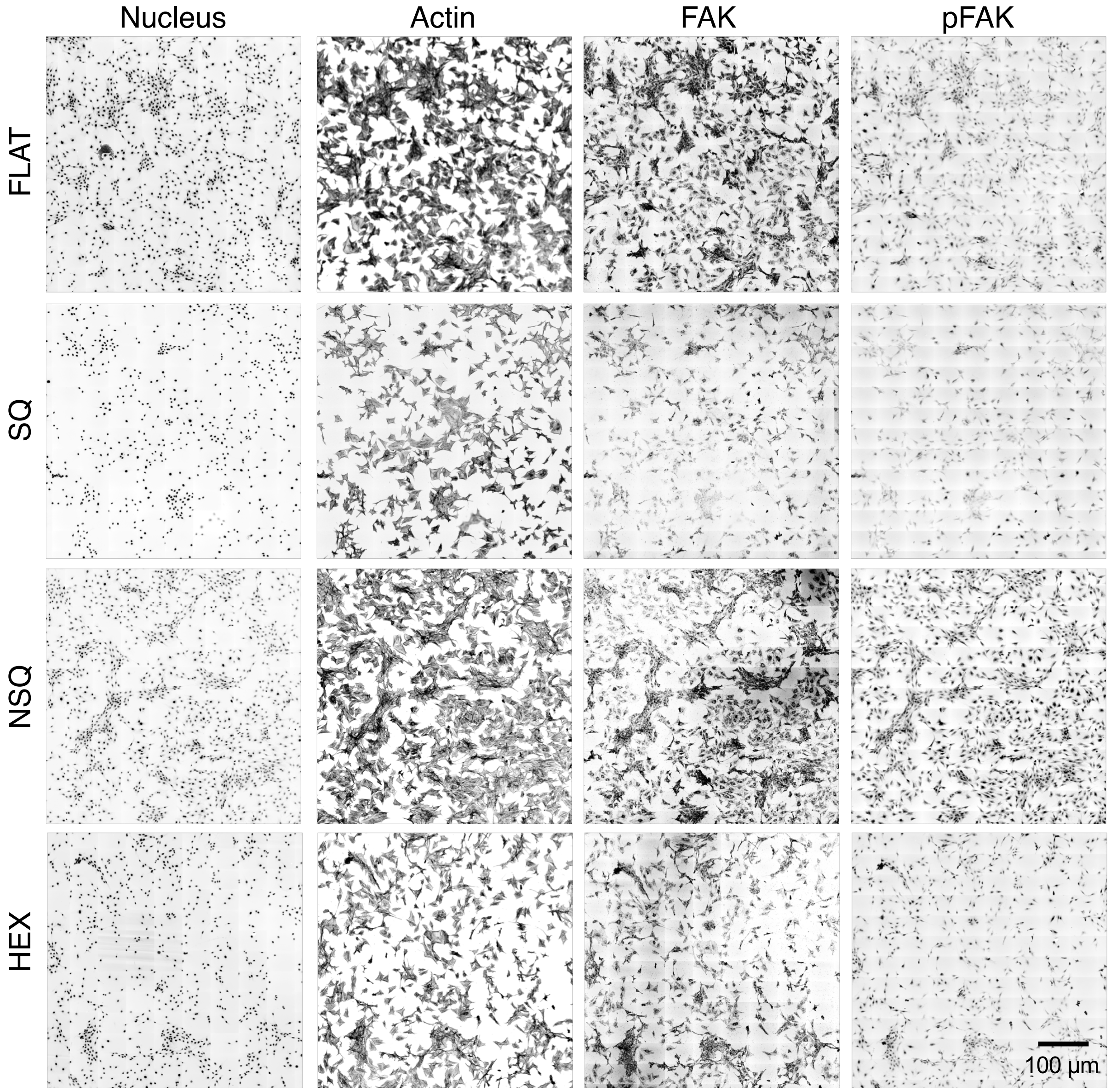


Figure S12. Overview of co-culture of pre-osteoblasts and fibroblasts on entire nanotopographies. Images of cells were captured at day 2 using 40X magnification and stitched together to form a tiled image of the entire nanotopography, which was then used for morphome extraction and subsequent analysis. A 2.12 mm x 2.12 mm grid was imaged for each nanotopography. Overview shows homogeneous cell attachment across all nanotopographies. Scale bar = 100 µm.

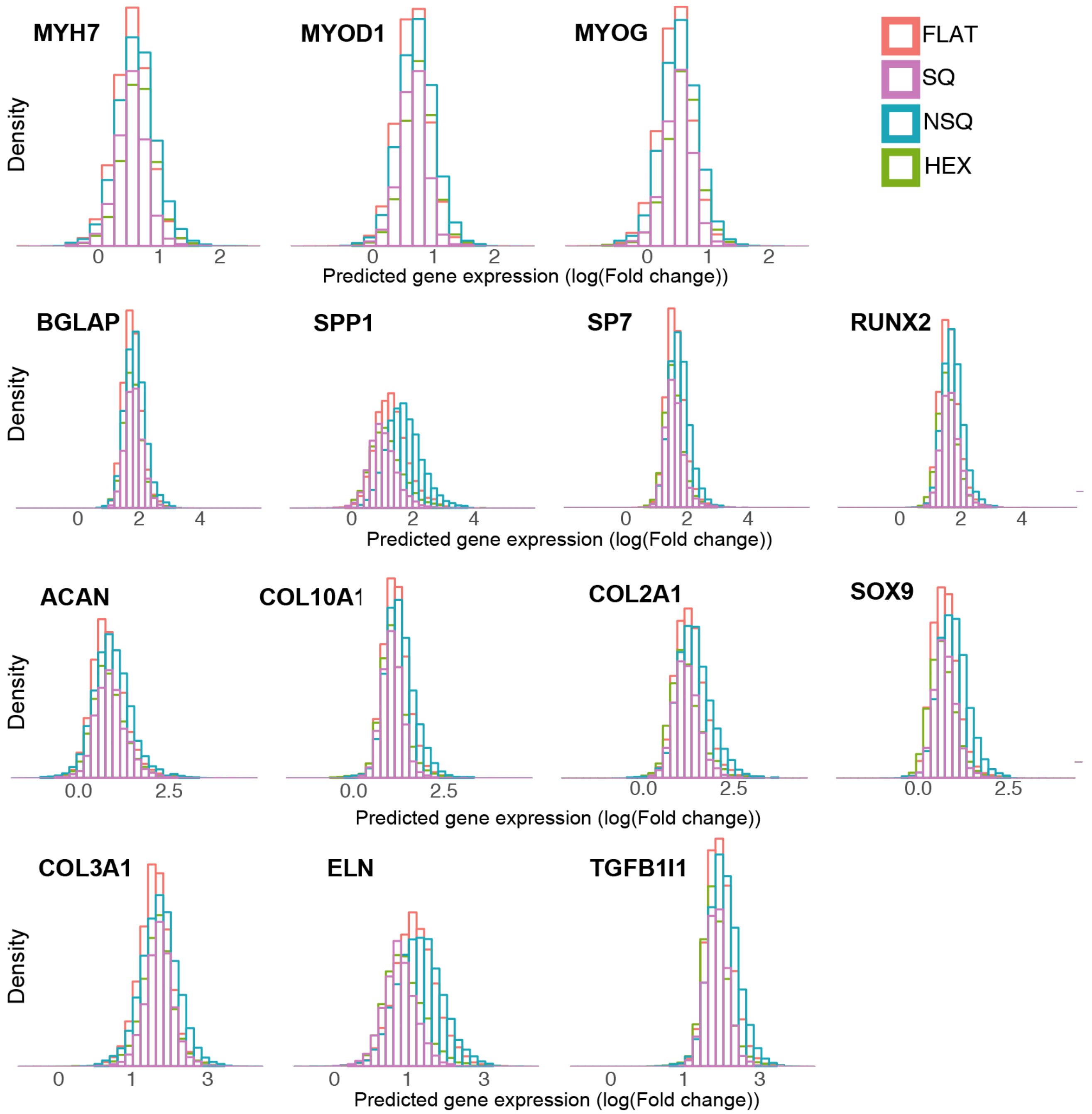


Figure S13. Histograms of predicted gene expression from pre-osteoblasts and fibroblasts co-cultured on nanotopographies. The morphome was extracted from all cells on the entire nanotopography. The morphome from the co-culture dataset was then centered to the mean and normalized to the standard deviation of the dataset used to create the predictive linear models. To predict the effect of nanotopographies on cell phenotype, the co-culture morphome was then used as input into the Bayesian linear regression model to obtain qPCR values for 14 different genes.

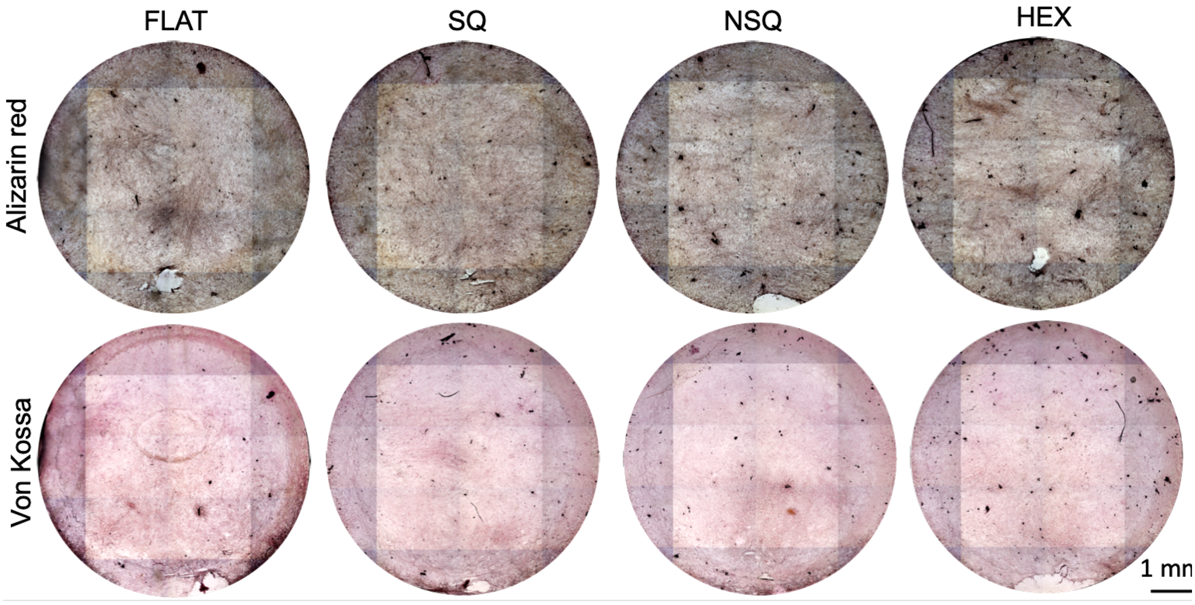


Figure S14. Mineralization of cocultured pre-osteoblasts and fibroblasts.

(A)

(B)

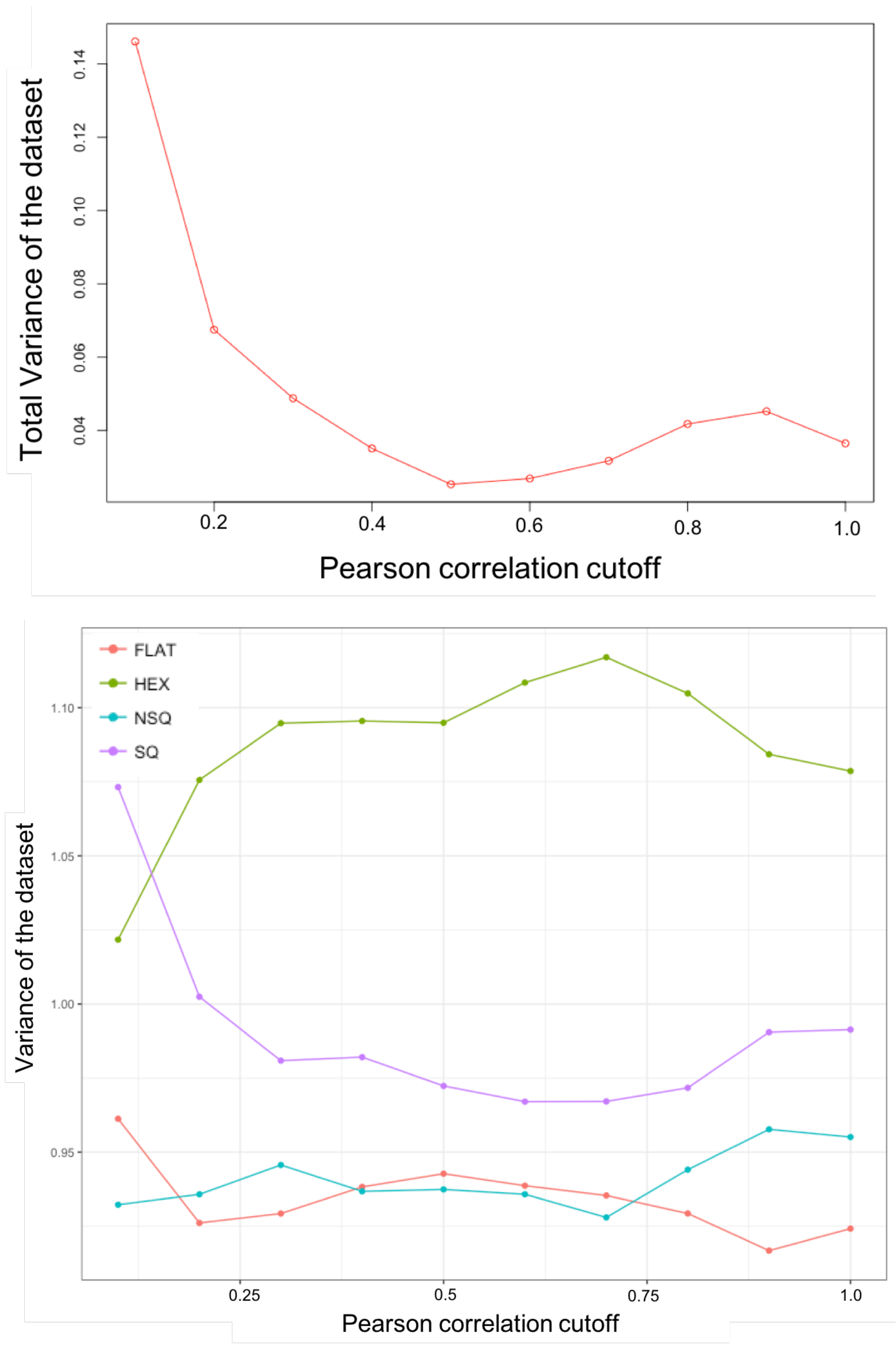


Figure S15. Effect of using different Pearson correlation cutoffs on the (A) total variance of the data set or (B) the variance of the topography-specific dataset. At a correlation cutoff of 0.9, the data set considerably reduced in feature number without compromising on the dataset variance.
